# A novel mouse model based on intersectional genetics enables unambiguous *in vivo* discrimination between plasmacytoid and other dendritic cells and their comparative characterization

**DOI:** 10.1101/2022.01.07.475382

**Authors:** Michael Valente, Nils Collinet, Thien-Phong Vu Manh, Karima Naciri, Gilles Bessou, Laurine Gil, Pierre Milpied, Elena Tomasello, Marc Dalod

**Author notes:** Co-senior authors (Elena Tomasello, Marc Dalod).

## Abstract

Plasmacytoid dendritic cells (pDC) were identified about 20 years ago, based on their unique ability to rapidly produce copious amounts of all subsets of type I and type III interferon (IFN-I/III) upon virus sensing, while being refractory to infection. Yet, the identity and physiological functions of pDC are still a matter of debate, in a large part due to their lack of specific expression of any single cell surface marker or gene that would allow to track them in tissues and to target them *in vivo* with high specificity and penetrance. Indeed, recent studies showed that previous methods that were used to identify or deplete pDC also targeted other cell types, including pDC-like cells and transitional DC (tDC) that were proposed to be responsible for all the antigen presentation ability previously attributed to steady state pDC. Hence, improving our understanding of the nature and *in vivo* choreography of pDC physiological functions requires the development of novel tools to unambiguously identify and track these cells, including in comparison to pDC-like cells and tDC. Here, we report successful generation of a pDC-reporter mouse model, by using an intersectional genetic strategy based on the unique co-expression of *Siglech* and *Pacsin1* in pDC. This pDC-Tomato mouse strain allows specific *ex vivo* and *in situ* detection of pDC. Breeding them with *Zbtb46^GFP^* mice allowed side-by-side purification and transcriptional profiling by single cell RNA sequencing of *bona fide* pDC, pDC-like cells and tDC, in comparison to type 1 and 2 conventional DC (cDC1 and cDC2), both at steady state and during a viral infection, revealing diverging activation patterns of pDC-like cells and tDC. Finally, by breeding pDC-Tomato mice with *Ifnb1^EYFP^* mice, we determined the choreography of pDC recruitment to the micro-anatomical sites of viral replication in the spleen, with initially similar but later divergent behaviors of the pDC that engaged or not into IFN-I production. Our novel pDC-Tomato mouse model, and newly identified gene modules specific to combinations of DC types and activations states, will constitute valuable resources for a deeper understanding of the functional division of labor between DC types and its molecular regulation at homeostasis and during viral infections.

## Introduction

Host survival to viral infections critically depends on type I interferons (IFN-I), which are cytokines endowed both with a direct and strong antiviral activity and with immunoregulatory functions^1^. The receptor for IFN-I is ubiquitously expressed and its triggering induces the expression of hundreds of genes, called interferon-induced genes (ISG). Many ISG encode viral sensors or viral restriction factors. Hence, IFN-I directly promote viral control by boosting intrinsic antiviral immunity in virtually all of the cells in the body^1^. IFN-I also promote the activation of immune cells^1^; they contribute to the maturation of dendritic cells (DC) that is necessary for the activation of antiviral immune effector cells including natural killer and CD8 T cells. IFN-I can also directly promote the activation of T cells and the production of antibodies by B cells^1^. However, excessive or dysregulated production of IFN-I can contribute to severe immunopathology, as observed in a number of autoimmune or inflammatory diseases, as well as in certain viral infections, including those by human immunodeficiency virus type 1 or by certain strains of Influenza viruses^1^. The role of IFN-I in the current Covid-19 pandemic caused by infection with SARS-Cov2 is complex, with harmful effects not only of failure to produce IFN-I, but also of delayed/chronic IFN-I production^2, 3^. Hence, understanding the cellular sources of IFN-I during viral infections and their molecular regulation is extremely important to be able to design treatments to boost or on the contrary to dampen this immune response, to promote health over disease depending on the pathophysiological context.

Plasmacytoid dendritic cells (pDC) are innate immune cells specialized in the rapid production of high levels of all types of IFN-I in response to many viruses^1, 4^. This process is triggered by the engulfment in pDC of exogenous virus-derived material that is routed into dedicated endosomes equipped with sensors specialized in the recognition of nucleic acids, namely Toll-like receptor 9 (TLR9) for unmethylated CpG DNA and TLR7 for single-stranded RNA. TLR7 and TLR9 then activate a MyD88-to-IRF7 signaling cascade leading to IFN-I production. However, only a very small fraction of pDC is activated for IFN-I production during viral infections *in vivo*^5^ and we still lack important information on how this process is regulated. In particular, we do not know precisely when, where and how pDC sense viral infection and sample viral-derived material, whether access to viral material is a key limiting step for pDC IFN-I production, and how pDC IFN-I production shapes host antiviral defense^4^. Answering these questions has been hampered by the lack of mutant mouse models enabling specific and penetrant targeting of pDC^6^, to locate them *in situ* in infected tissues and determine the nature and choreography of their interactions with infected cells or other cell types, to deplete them, or to interrogate their molecular regulation through *in vivo* gene editing. This bottleneck was largely caused by the fact that no gene is specifically expressed in pDC and not in other cell types, such that genetic targeting of pDC could not be achieved by classical knock-out or knock-in approaches for a single gene. This is all the more the case as, recently, novel DC types were identified that share cell surface marker and gene expression with pDC^7–9^, including pDC-like cells and transitional DC (tDC), which were often contaminating the cell populations studied as pDC, and which were perturbed by the treatments aimed at targeting pDC, in previous studies. Hence, it is not yet clear to which extent some of the properties attributed to pDC could have been due to their contamination by pDC-like cells or tDC, including cognate activation of T cells and the induction of immune tolerance^7–9^. Hence, there is an unmet scientific need to generate innovative mutant mouse models for specific and penetrant targeting of pDC^6^, as well as to further characterize pDC-like cells and tDC in comparison to pDC^9^, to help better discriminating these cell types from one another and to lay the foundations for future studies aiming at dissecting their functional specialization.

Here we report the successful design of a pDC-reporter mouse model, enabling unambiguous identification of pDC *ex vivo* by flow cytometry, and *in situ* in tissues by confocal microscopy. Moreover, we illustrate the usefulness of this novel mutant mouse model by harnessing it to determine the choreography of pDC recruitment to the micro-anatomical sites of viral infection in the spleen, before their production of IFN-I, and the later differential behavior of these cells depending on their activation state. We also used this novel mouse model to unambiguously isolate pDC, pDC-like cells and tDC at steady state and during the infection, to compare their gene expression profiles by single cell RNA sequencing (scRNA-seq), with inclusion of type 1 and 2 conventional DC (cDC1 and cDC2) as key reference DC types. Our novel pDC-reporter mouse model and the associated scRNA-seq dataset will constitute precious resources for future research on the *in vivo* functions of DC types and their molecular regulation.

## Results

### pDC-Tomato mice allow specific detection of pDC in whole body by fluorescence

The generation of mice specifically targeting pDC has been hampered by the lack of any molecule expressed only by pDC, which was confirmed when we attempted to generate pDC reporter mice by targeting the *Siglech* gene, whose product is highly expressed by mouse pDC^10, 11^. Indeed, fate-mapping of *Siglech*-targeted cells, using *Siglech^iCre^* knock-in mice bred with *Rosa26^LoxP-STOP-LoxP(LSL)^*^-*RFP*^ mice^12^ (Figure 1A), showed that over 95% of splenic pDC were RFP^+^ (Figure 1B and S1A-B), but variable proportions of myeloid and lymphoid lineages also expressed RFP (Figure 1B and S1A-B). Hence, although this strategy enabled extremely efficient targeting of pDC, it was not specific enough, confirming that *Siglech* is expressed in a fraction of conventional dendritic cells (cDC) or their immediate precursors^13^ and in other myeloid cells^14^.

**Figure 1.**
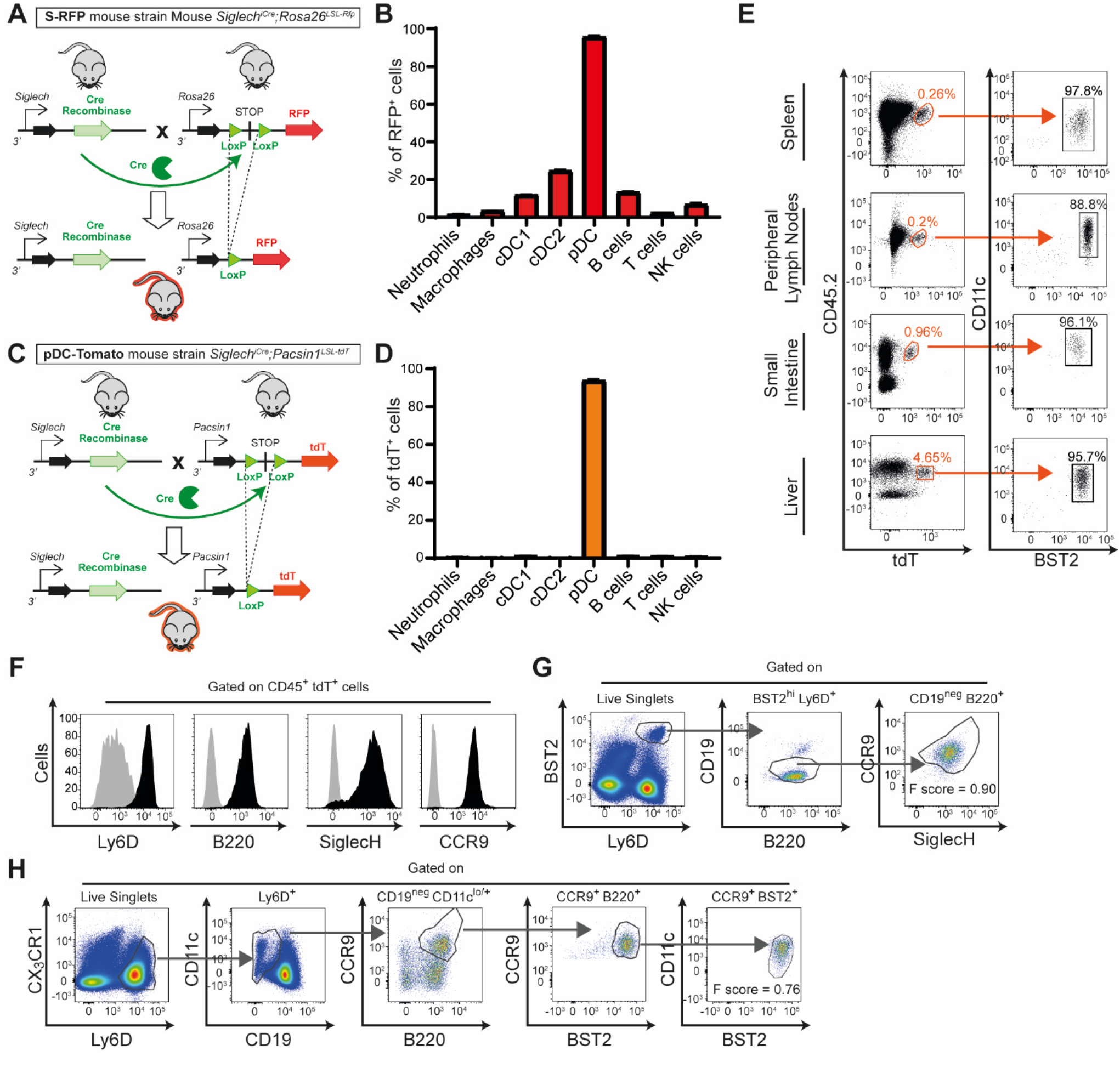
pDC-Tomato mice allow specific and unambiguous identification of pDC in different organs. (A) Scheme illustrating the strategy followed to generate S-RFP mice. LoxP is the sequence recognized by the Cre recombinase. STOP corresponds to a transcriptional stop sequence. (B) Splenocytes isolated from S-RFP mice were stained with fluorescently labeled antibodies in order to identify the indicated myeloid and lymphoid cell populations and analyze their expression of RFP by flow cytometry. The data shown are pooled from 2 independent experiments, with 8 animals in total. (C) Scheme illustrating the strategy followed to generate pDC-Tomato mice. (D) Splenocytes isolated from pDC-Tomato mice were stained as in (B) to analyze their expression of tdT by flow cytometry. The data shown are pooled from 2 independent experiments, with 6 animals in total. (E) Single cell suspensions of different organs from pDC-Tomato mice were stained with indicated fluorescent antibodies and analyzed by flow cytometry. (F) Splenocytes from (E) were analyzed for the expression of indicated markers on CD45^+^ tdT^+^ cells. Grey histograms correspond to negative controls (fluorescent minus one). Black histograms correspond to the signal obtained upon staining with the indicated antibody. For (E-F), the data shown are from one mouse representative of 7 animals for spleen, and of 5 animals for peripheral lymph nodes (LN), liver and small intestine. (G-H) The HyperFinder plugin of the Flowjo software was applied to define an unsupervised gating strategy to identify pDC from uninfected (G) or 36h-MCMV-infected (H) pDC-Tomato mice.

We then reasoned that specific targeting of pDC would be achievable by harnessing intersectional genetics, namely by driving expression of the fluorescent reporter under the control of two genes known to be co-expressed only in pDC. Specifically, we hypothesized that this should be possible if *Siglech*-driven Cre-recombinase activity activates a conditional fluorescent reporter cassette knocked in the 3’-UTR of a gene exclusively expressed by pDC within *Siglech* fate-mapped cells. For this purpose, we selected *Pacsin1,* a gene expressed exclusively in pDC within hematopoietic cells^15^ and coding for a molecule involved in the regulation of TLR7/9-dependent IFN-I production by pDC^16^. We thus generated *Pacsin1^LoxP-STOP-LoxP-tdTomato^* (*Pacsin1^LSL-tdT^*) mice and crossed them with *Siglech*^iCre^ mice, thereby generating pDC-Tomato mice (Figure 1C). In splenocytes from pDC-Tomato mice, tdT was exclusively expressed in pDC, without any specific signal detectable in the other myeloid and lymphoid cells analysed (Figures 1D and S1A,C). tdT^+^ cells did not express lineage markers or CD11b (Figure S1D). Moreover, the CD45^+^ tdT^+^ cells isolated from different lymphoid (spleen, lymph nodes) and non-lymphoid organs (intestine, liver) of pDC-Tomato mice uniformly expressed intermediate levels of CD11c and high levels of BST2 (Figure 1E), strongly suggesting that they were all pDC^17^. Indeed, CD45^+^ tdT^+^ cells homogenously co-expressed several markers whose combination is specific to pDC, namely Ly6D, B220, SiglecH and CCR9 (Figure 1F). Altogether, these results show that tdT expression in pDC-Tomato mice is sufficient *per se* to specifically and unambiguously identify pDC, thus validating our intersectional genetics strategy for specific pDC targeting based on their unique co-expression of *Siglech* and *Pacsin1*.

### pDC-Tomato mice allowed designing a novel gating strategy to unambiguously identify pDC in homeostatic and inflammatory conditions based only on cell surface marker expression

The phenotypic definition of pDC is complex. No single cell marker allows their specific identification. Different research teams use different marker combinations to achieve this aim. Attempting to define pDC as cells co-expressing CD11c, BST2 and SiglecH can lead to their contamination by cDC, pDC-like cells^7^ or tDC^8, 9, 18^, depending on the precise threshold used for individual marker expression levels. This issue is made further complex under inflammatory conditions, since they alter the expression of these markers. For example, during systemic infection by mouse cytomegalovirus (MCMV), CD11c expression levels are increased on pDC and decreased on activated cDC, whereas SiglecH expression is decreased on pDC and BST2 increased on cDC, causing a convergence of the phenotype of pDC and cDC^5^. Hence, we took advantage of the specificity of our novel pDC-Tomato mouse model to seek for an adequate gating strategy allowing unequivocal pDC identification both at steady state and during MCMV infection. To this aim, we defined *bona fide* pDC as tdT^+^ cells and used the HyperFinder algorithm to identify an unsupervised gating strategy enabling the identification of these cells based exclusively on combinations of cell surface markers, without using their expression of tdT. At steady state, splenic pDC were identified as Bst2^high^ Ly6D^+^ cells, positive for B220 but negative for CD19, and homogenously expressing CCR9 and SiglecH (Figure 1G). During MCMV infection, splenic pDC were identified as Ly6D^+^ cells with varying but not very high levels of CX_3_CR1, positive for CD11c but negative for CD19, and homogenously expressing high levels of CCR9, B220 and BST2 (Figure 1H). Taken together, these results indicate that current identification of pDC as lin^neg^ CD11b^neg^ CD11c^low-to-int^ BST2^high^ cells^17^ can be improved by addition of positivity for Ly6D or CCR9 and eventually exclusion of CX_3_CR1^high^ cells. We thus propose to unambiguously identify pDC in homeostatic and inflammatory conditions as Ly6D^+^ BST2^high^ CD19^neg^ B220^+^ CD11b^neg^ CD11c^+^ cells.

### In pDC-Tomato mice, tdT starts to be expressed exclusively in late stages of pDC differentiation

We next analysed bone marrow cells to investigate the expression of tdT along the myeloid^19–22^ and lymphoid^7, 8^ ontogeny paths that have been proposed for pDC (Figure 2A). Within differentiated bone marrow cells, tdT was expressed exclusively in pDC (Figure 2B); it was undetectable in other differentiated immune cell types including cDC, consistent with the results observed in other organs (Figure 1). tdT could also be clearly detected, although at a lower mean fluorescence intensity, in the immediate precursors of pDC, the pre-pDC (Figure 2C), defined as Lin^neg^ B220^+^ CD11c^+^ MHC-II^neg^ FLT3^+^ BST2^high^ SiglecH^+^ (Figure S2A). Upstream along the proposed myeloid path of pDC differentiation^13, 19–22^, very low levels of tdT could be detected in SiglecH^+^ pre-DC, but not in SiglecH^neg^ pre-DC, irrespective of their expression of Ly6C (Figure 2D and Figure S2A). This is consistent with SiglecH^+^Ly6C^+/neg^ pre-DC giving rise to both cDC and pDC, whereas SiglecH^neg^Ly6C^+^ and SiglecH^neg^Ly6C^neg^ generate mostly cDC2 versus cDC1 with only few pDC, respectively^13^. Further upstream, the common DC progenitor (CDP), monocyte and DC progenitor (MDP) and common myeloid progenitor (CMP) were negative for tdT expression (Figure 2E and Figure S2A).

**Figure 2.**
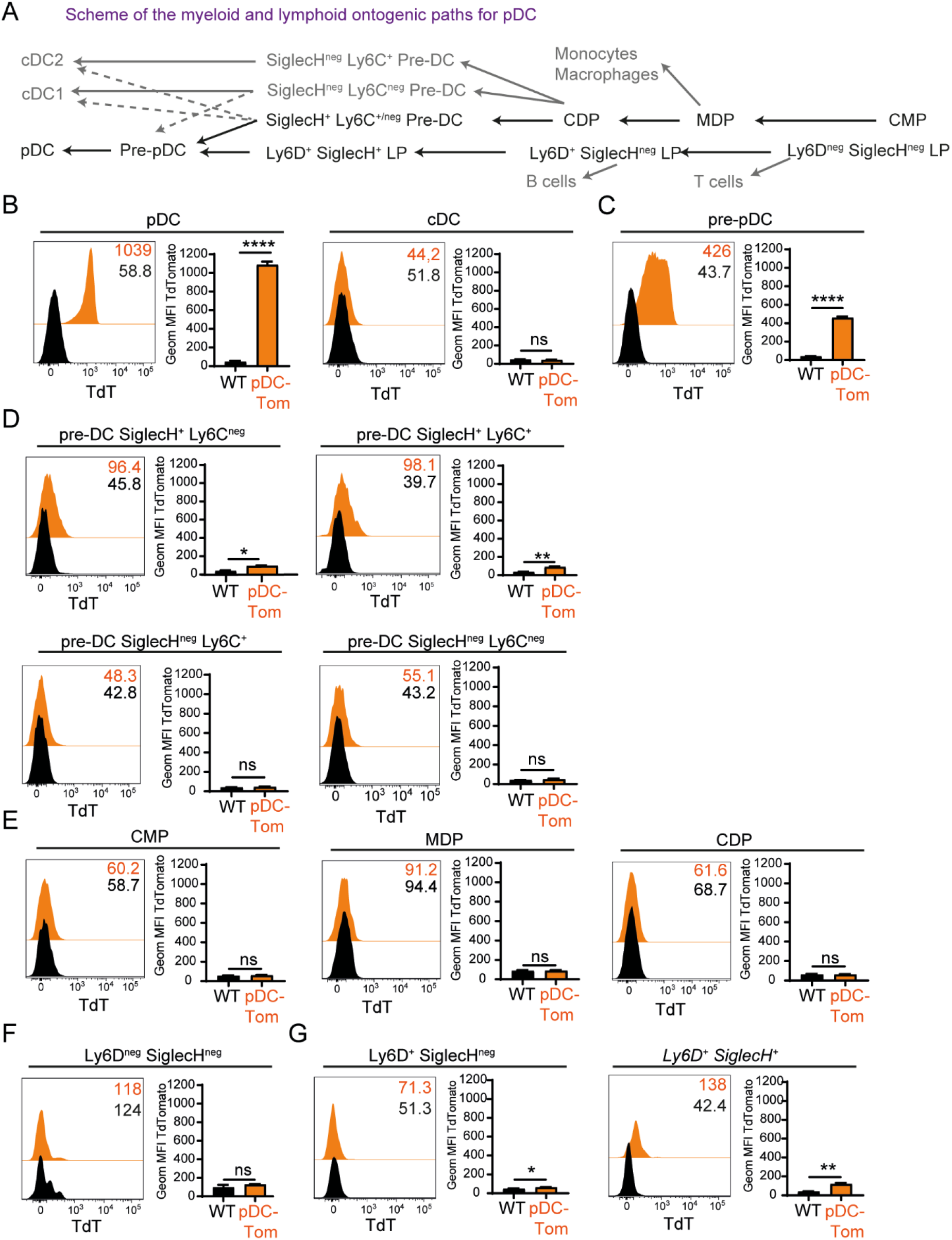
tdT expression is detectable in late bone marrow precursors selectively committed to the pDC lineage. (A) Scheme of the previously proposed ontogenic paths for pDC differentiation along the myeloid (top) or lymphoid (bottom) lineages. (B-G) Bone marrow cells were isolated from pDC-Tomato animals, stained with fluorescently labeled antibodies and analyzed by flow cytometry. The expression of tdT (orange histograms) was evaluated in BM pDC (and cDC (B), pre-pDC (C) and in different cell types along the myeloid (D-E) or lymphoid (F-G) ontogenic paths. (B-G) Wt C57BL/6 mice were used as negative controls (black histograms). The fluorescence histograms shown (left) are from on mouse representative of 5 pDC-tomato animals from 2 independent experiments. The bar graphs (right) correspond to the results of all 5 animals, with data shown as mean ± SEM. Statistical analysis by unpaired t test, \**p <* 0.05, \*\**p <* 0.01, \*\*\**p <* 0.001, \*\*\*\**p <* 0.0001.

Upstream along the proposed lymphoid path of pDC differentiation, low but clearly detectable levels of tdT could be detected in CD127^+^ SiglecH^+^ Ly6D^+^ progenitors (Figure 2F and Figure S2B), consistent with these cells giving rise specifically and efficiently to pDC^7^. Very low levels of tdT could be detected in the upstream Ly6D^+^ SiglecH^neg^ lymphoid progenitor, but not anymore further upstream in the Ly6D^neg^ SiglecH^neg^ lymphoid progenitor (Figure 2G and Figure S2B).

Taken together, our results show that, in pDC-Tomato mice, tdT expression is exclusively induced in late bone marrow precursors already committed to the pDC lineage, with a strong increase of expression in pre-pDC and maximal expression in differentiated pDC.

### ZeST mice allow discriminating *bona fide* pDC from pDC-like cells and tDCs

We next wanted to confirm that tdT expression in pDC-Tomato mice enabled discriminating *bona fide* pDC from pDC-like cells^7^ and tDC^8, 9, 18^. Although refined phenotypic keys have been proposed to discriminate *bona fide* pDC from pDC-like cells and tDC by flow cytometry at steady state^9^, there is still an unmet need for reporter mouse models allowing their specific tracing under inflammatory conditions as well as *in situ* in tissues by immunohistofluorescence. Within hematopoietic cells, the *Zbtb46* gene is specifically expressed in the cDC lineage^15^, including in pre-cDC^23^ and in pDC-like cells^7^ and tDC^9^. This has been confirmed previously in pre-cDC and pDC-like cells by using the *Zbtb46*^GFP^ reporter mice^7, 23^. Therefore, to rigorously identify pDC-like cells and tDC in our pDC-Tomato mice, we crossed them with *Zbtb46*^GFP^ reporter mice to generate ***Z****btb46*^GFP^;***S****iglech*^iCre^;*Pacsin1^LSL-^****^t^****^dT^* animals, hereafter called ZeST mice (Figure 3A).

**Figure 3.**
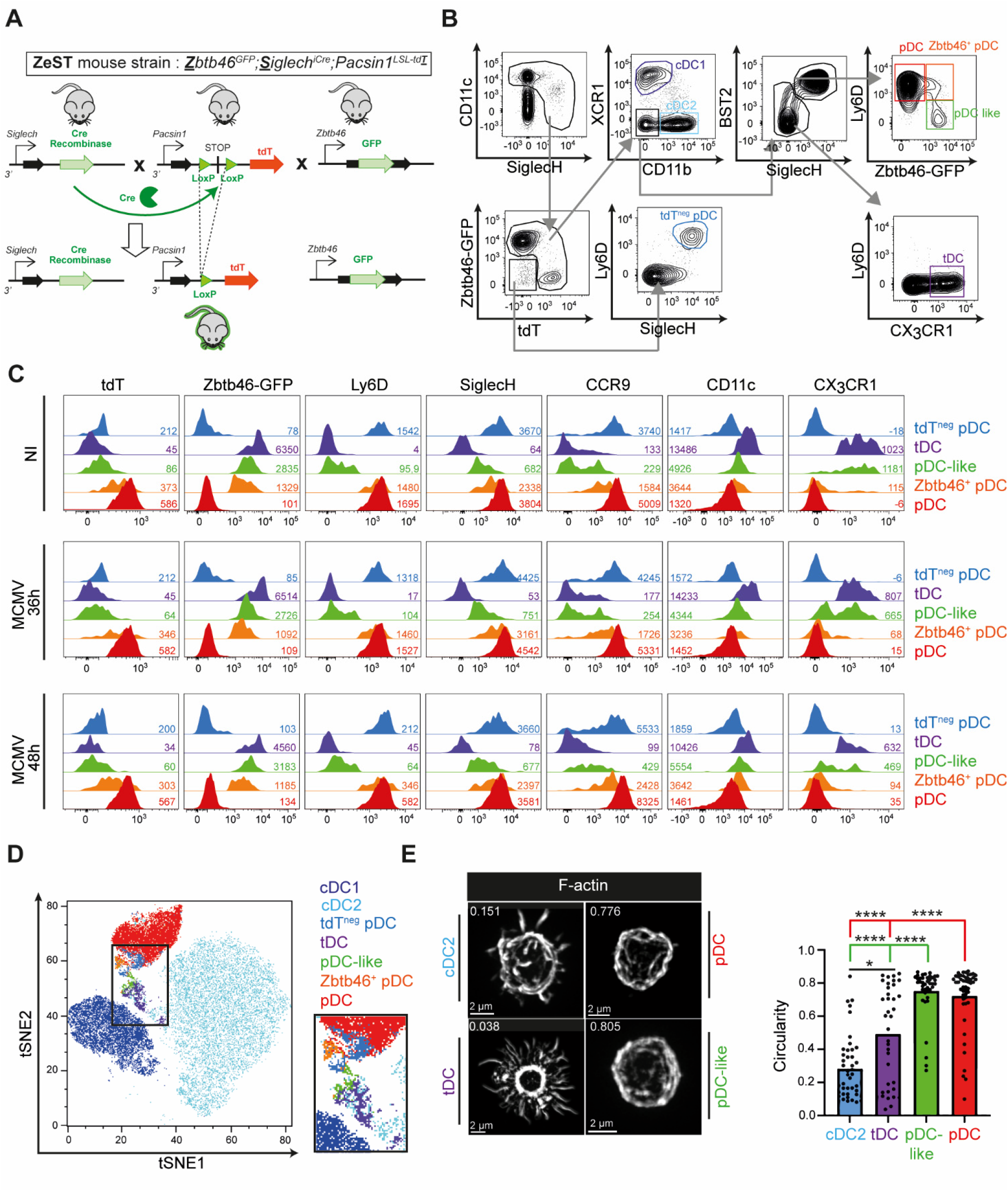
ZeST mice allows to unambiguously discriminate pDC from tDC and pDC-like cells. (A) Scheme illustrating the strategy followed to generate ZeST mice. (B-C) Splenocytes from ZeST mice were stained with fluorescently labeled antibodies and analyzed by flow cytometry. (B) Gating strategy illustrating the phenotypic criteria followed to identify cDC1, cDC2, tDC, pDC like cells, pDC, Zbtb46^+^ pDC and tdT^-^ pDC. Autofluorescent, Lineage^+^ and dead cells were excluded at the beginning of the strategy. (C) Expression of indicated fluorescent proteins or cell surface markers on each of the cell populations gated as in (B), from spenocyte suspensions from uninfected vs 36h or 48h MCMV-infected pDC-Tomato mice. (D) Projection of the cell types identified in (B), on the tSNE space calculated with the Flowjo plugin for all cells from the gate encompassing all cells expressing high levels of CD11c or positive for SiglecH. Red, pDC; medium blue, tdT^neg^ pDC; orange, Zbtb46^+^ pDC; green, pDC-like cells; purple, tDC. The data shown are from one ZeST mouse representative of at least 10 uninfected animals for (B-D), and for 7 MCMV-infected animals at 36h p.i or 8 MCMV-infected animals at 48h p.i. for (C). (E) Quantitative and unbiased assessment of the cellular morphology of cDC2, tDC, pDC-like cells and pDC sorted from the spleen of uninfected ZeST mice according to the gating strategy shown in (B). One representative confocal microscopy image of each DC type is shown on the left. The distribution of the circularity indices for individual cells across DC types is shown as dots on the right, with the overlaid color bars showing the mean circularity indices of each DC type. The data shown are from 2 independent experiments each performed with one mouse, with 37 to 44 individual cells analyzed for each DC type. Kruskal-Wallis was used for the statistical analysis, *P<0.05, **P<0.01, ***P<0.001, ****P<0.0001.

In ZeST mice, most of the lin^neg^ cells expressing high levels of CD11c or being positive for SiglecH expressed either GFP or tdT, in an almost mutually exclusive manner (Figure 3B). Within cells expressing tdT or GFP, we identified cDC1 as XCR1^+^ and cDC2 as CD11b^+^ (Figure 3B). Within the XCR1^neg^ CD11b^neg^ BST2^high^ SiglecH^+^ gate, we identified pDC as Ly6D^+^ cells, with only a small fraction expressing GFP (Zbtb46^+^ pDC); pDC-like cells were identified as Ly6D^neg^ GFP^+7^. Within the XCR1^neg^ CD11b^neg^ BST2^neg-to-int^ SiglecH^neg^ gate, we identified tDC as Ly6D^neg^ CX_3_CR1^+^ cells^9^ (Figure 3B). Only very few of the Lin^neg^, CD11c^high^ or SiglecH^+^, cells expressed neither GFP nor tdT; yet, a fraction of these cells expressed high levels of both Ly6D and SiglecH, suggesting that they were tdT^neg^ pDC (Figure 3B).

To validate the identity of all the cell types identified by manual gating, we further characterized their phenotype (Figure 3C). As expected, beyond expressing high levels of Ly6D and being SiglecH^+^, all pDC populations strongly expressed CCR9 and tdT, while harboring intermediate levels of CD11c. Beyond expressing CX_3_CR1 but not Ly6D, tDC expressed high levels of GFP and were negative for tdT. Beyond expressing SiglecH, BST2 and GFP but not Ly6D, pDC-like cells did not express tdT and no or low levels of CCR9. Hence, as expected, tdT was only expressed in cells harbouring a phenotype of *bona fide* pDC, whereas GFP was mainly expressed in cDC1, cDC2, pDC-like cells and tDC. The proportions of these distinct populations, as well as their surface phenotype, were not significantly modified during MCMV infection (Figures 3C and S3A-I).

We performed an unsupervised analysis of the phenotypic relationships between all the Lin^neg^, CD11c^high^ or SiglecH^+^, cells by generating a tSNE bi-dimensional representation of the data with FlowJo (Figure 3D). We then projected on the tSNE space the populations identified through manual gating as defined in panel B. This analysis identified three main clusters of cells, corresponding to cDC2 (pale blue), cDC1 (dark blue) and pDC (red). The small populations of tdT^neg^ pDC (intermediate blue) and Zbtb46^+^ pDC (orange) were found in tight proximity to the pDC cluster, consistent with their pDC identity. The pDC-like cells (green) and tDC (purple) settled between the pDC and cDC2 clusters, consistent with previous similar observations^9^.

Finally, we sorted cDC2, tDC, pDC-like cells and pDC from steady state mouse spleens and examined their morphology using confocal microscopy (Figure 3E). cDC2 harboured irregular morphologies with many pseudopods or dendrites, translating into a low circularity index. In contrast, the overwhelming majority of pDC-like cells and pDC harboured a rather round morphology, translating into a high circularity index. The tDC population presented a bimodal distribution of circularity indices, with about half of the cells harbouring a dendritic morphology akin to that of cDC2, and the other half a round morphology akin to that of pDC. Hence, quantitative and unbiased analysis of cellular morphology was consistent with success in high degree purification of the different DC types examined when using ZeST mice, but also highlighted the heterogeneity of the tDC population, consistent with its inclusion of two subsets based on their expression pattern of a limited selection of fingerprint genes and cell surface proteins: an Ly6C^int/high^ CD11c^low/int^ MHC-II^neg/low^ pDC-like subset, and a Ly6C^neg/low^ CD11c^high^ MHC-II^int/high^ cDC-like subset^9^.

Altogether, these results confirmed that ZeST mice allowed properly discriminating pDC from pDC-like cells and tDC. Yet, they also unravelled rare fractions of putative pDC devoid of tdT or expressing *Zbtb46*.

### scRNA-seq confirmed proper identification of pDC, pDC-like cells and tDC in ZeST mice, and enabled refining their transcriptomic characterization

Next, we took advantage of the ZeST mice to perform single cell gene expression profiling of all the five DC types, upon index sorting from the spleen of uninfected animals and of mice that had been infected with MCMV for 36h or 48h, following the gating strategy presented in the previous section, and using the FB5P-seq protocol^24, 25^. We first focused on the analysis of the cells isolated at steady state, to generate DC type-specific signatures to be used for the annotation of the cell clusters of the whole dataset. Following classical technical quality controls, we retrieved 343 cells that were clustered by Seurat based on their gene expression profiles (Figure S4A) and by R-phenograph based on their cell surface marker expression as assessed from the index sorting flow cytometry data, without taking tdT expression into account (Figure S4B). The 7 clusters were annotated to the DC types based on the expression pattern of archetypic cell surface markers for R-phenograph (Figure S4C) and on their marker genes for Seurat. The consistency between the annotations of individual cells as belonging to one of the five DC types based on phenotype (R-Phenograph) versus gene expression (Seurat) was very good (Figure S4D). However, Seurat cluster 1 appeared to encompass a mixture of tDC and pDC-like cells, and Seurat cluster 2 of tDC and cDC2, consistent with the known proximity between these DC types, with tDC (Lin^neg^ CD11c^+^ CD11b^neg^ CX_3_CR1^+^ cells) reported to encompass two subpopulations, one Ly6C^int/high^ CD11c^int^ closer to pDC and the other Ly6C^neg/low^ CD11c^high^ closer to cDC2^9^. Therefore, to identify in a robust and unbiased manner the individual cells that we could unambiguously assign to each of the DC type, we used a combinatorial strategy whereby, to be selected, a cell had to belong to the expected intersection between the R-Phenograph and Seurat clusters (green cells in Figure S4D) and to be enriched for the corresponding DC type-specific transcriptomic signatures, as established from independent datasets^7, 26^ and tested by using the sgCMAP algorithm^27^ (Figure S4E). This led to a robust final cell type assignment for 205 cells, namely 103 pDC, 26 pDC-like, 19 tDC, 23 cDC2 and 34 cDC1 (Figure S4E-F). We then generated cell type-specific transcriptomic signatures from this dataset, to corroborate assignment of DC types to the 951 cells from the whole dataset that had passed both quality control and removal of other contaminating cell types (Figure S5A). In brief, following cell clustering with Seurat (Fig. S5A) and R-Phenograph (Figure S5B-C), we assigned a DC type to 851 individual cells based on their belonging to the expected intersection between these two clustering strategies (Figure S5D). Clustering the 951 cells based on their sgCMAP scores largely confirmed this final cell type assignment (Figure S5E). Importantly, comparing the sorting phenotype of the cells to their final cell type assignment showed that all the cells sorted as pDC remained *in fine* assigned to pDC when using this unbiased classification scheme (Fig S5E). This was also the case for the vast majority of the cells sorted as tdT^neg^ pDC. The assignment of individual cells to cDC1 was also consistent between cell sorting and the *a posteriori* re-annotation. Clustering of the cells on the sgCMAP scores also tended to confirm the distinction between cDC2 and tDC, although many cells had a null score for the “tDC_vs_cDC2” sgCMAP signature, emphasizing the proximity between these two DC types. In contrast, many cells sorted as Zbtb46^+^ pDC were pDC-like cells. In addition, some cells sorted as pDC-like were *in fine* assigned to the tDC type. This analysis led to a robust final cell type assignment for 851 cells, namely 310 pDC, 170 pDC-like, 146 tDC, 103 cDC2 and 122 cDC1, with 100 cells left unannotated (Figure 4A). These cells expressed the expected pattern of cell surface markers (Figure 4B). Of note, both pDC-like cells and tDC expressed higher levels of CX_3_CR1 than the other DC types, and lacked CD11b. However, akin to cDC2 (and cDC1), tDC expressed high levels of CD11c and GFP, whereas pDC-like cells expressed SiglecH, BST2, Ly6D and CCR9 to levels intermediate between those of pDC (high) and of cDC (low). As expected, only pDC expressed the signature genes *Ccr9*, *Klk1* and *Cox6a2*, as well as rearranged immunoglobulin genes (Figure 4C). Only pDC expressed tdT above autofluorescence levels (Figure 4D). Of note, the fraction of tdT^neg^ pDC is largely over-represented in this figure as compared to their proportions *in vivo* (see Figure 3B), because for the FB5P-seq experiment we strongly enriched these rare cells to be able to characterize them. Since we did not use tdT signal to analyse the FB5P-seq dataset, this result thus confirms the specificity of the pDC-Tomato reporter mouse for the identification of pDC. pDC and pDC-like cells shared high expression of the genes *Siglech* (Figures 4C, E), *Tcf4* and *Runx2* (Figure 4C). tDC shared with pDC-like cells high expression of the genes *Crip1*, *Lgals3* and *Vim* (Figure 4C), which have been previously reported to discriminate pDC-like cells from pDC^7^. pDC-like cells and tDC shared with cDC higher expression of the genes *Zbtb46*, *Spi1*, *Slamf7* and *S100a11*, as compared to pDC (Figure 4C). Contrary to cDC2, a fraction of pDC-like cells and tDC expressed the *Cd8a* gene (Figures 4C, E), as previously reported for tDC^9, 18, 26^. pDC-like cells, tDC and cDC2 specifically expressed the genes *Ms4a4c* (Figures 4C, E) and *Ms4a6c* (Figure 4C). tDC and cDC2 selectively shared expression of the genes *Ms4a6b* (Figure 4C, E) and *S100a4* (Figure 4C). tDC expressed higher levels of certain cDC genes than pDC and pDC-like, including *Batf3*, *Rogdi* and *Cyria* (Figure 4C). Finally, pDC-like cells expressed very high levels of *Ly6c2* (Figure 4C, E), consistent with the recent demonstration that Ly6C identifies a tDC subpopulation that resembles more pDC^9^, whereas high expression of CD11c identifies a tDC subpopulation resembling more cDC2^9^ as observed in our dataset for tDC when compared to pDC and pDC-like cells (Figure 4B). Hence, our results show that the pDC-like cells that we characterize here align with both the pDC-like cells initially described by the team of Roxane Tussiwand^7^, and with the fraction of the tDC population that resembles pDC as described by the teams of Juliana Idoyaga and Boris Reizis^9^, whereas the tDC that we characterize here correspond to the other fraction of the tDC population that resembles cDC2^9^. In conclusion, FB5P-seq confirmed proper identification of pDC, pDC-like cells and tDC in ZeST mice, and enabled refining their characterization by enabling for the first time a side-by-side transcriptomic comparison of these three cell populations, through whole genome expression profiling, at the single cell level, and with inclusion of cDC1 and cDC2 as key reference cell types.

**Figure 4.**
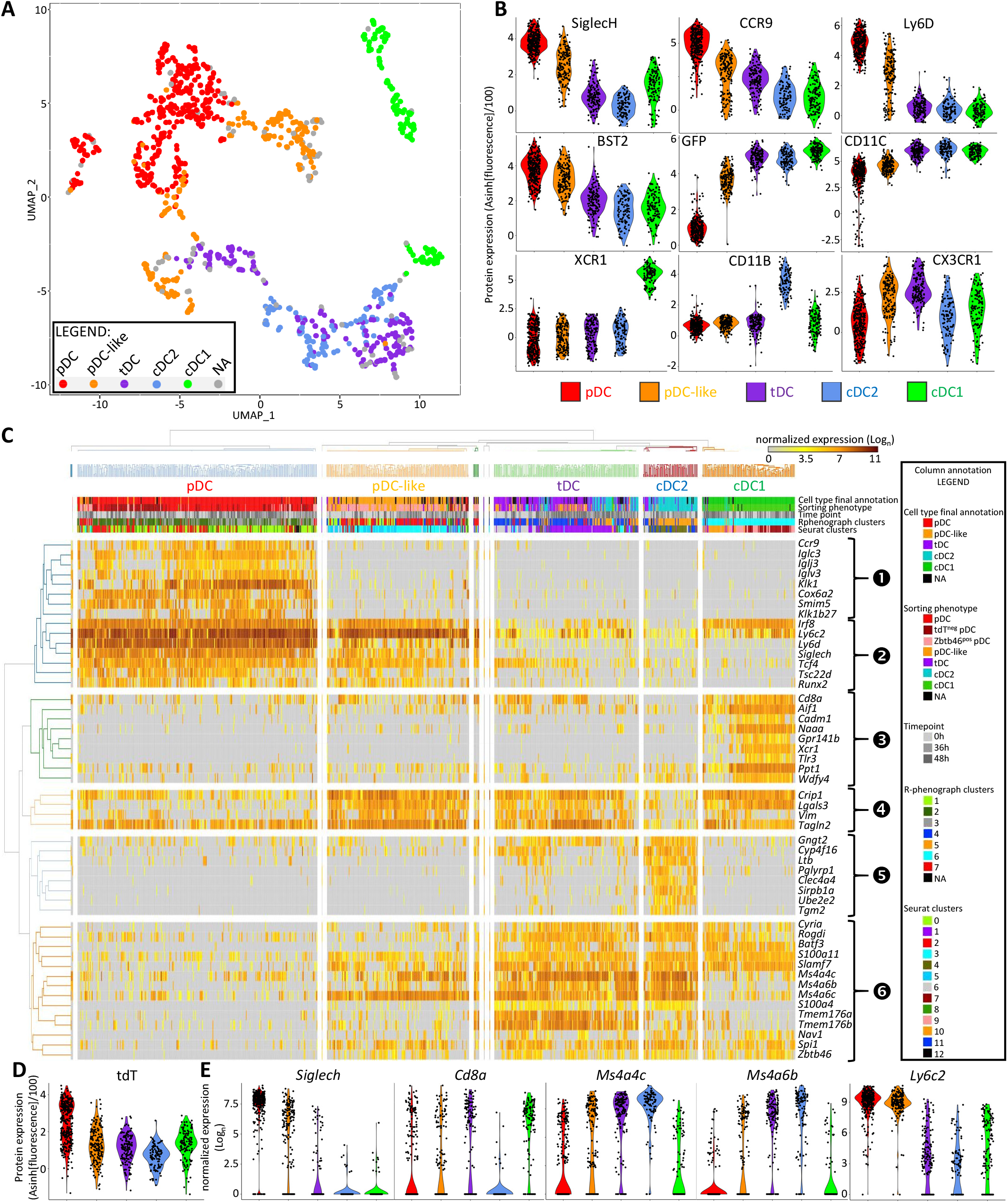
scRNA-seq confirmed proper identification of pDC, pDC-like cells and tDC in ZeST mice. (A) Dimensionality reduction performed using the UMAP algorithm for DC types isolated from the spleens of three ZeST (*Zbtb46^GFP^*;*Siglech^iCre^*;*Rosa26^LSL-tdT^*) mice (one uninfected, one MCMV-infected for 36h and the third infected for 48h). Cells were index sorted into the 5 DC types studied using the gating strategy presented in Figure 3B, and used for single cell RNA sequencing using the FB5P-seq procedure. Cells that did not pass the FB5P-seq quality controls or that turned out to be contaminants were removed from the analysis, leading to keeping 951 cells. As indicated by the color code, individual cells were re-assigned *a posteriori* a DC type identity (cDC1, cDC2, pDC, pDC-like or tDC), based on combined analysis of their phenotypic and transcriptomic characteristics, as assessed by R-phenograph clustering based on fluorescence signal intensities for 10 phenotypic markers (CD11c, XCR1, CD11b, GFP, SiglecH, BST2, Ly6D, CX_3_CR1 and B220) and by Seurat graph-based transcriptomic clustering with analysis of the marker genes of each cluster, with confirmation via a single cell cMAP-based enrichment analysis for DC type-specific signatures generated from prior analysis of the cells from uninfected mice only (see Figures S4 and S5), leading to 100 cells left non-annotated (NA). (B) Violin plots showing the expression of phenotypic markers across the DC types identified in (A). (C) Heatmap showing mRNA expression levels of selected genes (rows) across all the 851 individual cells annotated for DC types (columns), with hierarchical clustering of gene using Euclidian distance for genes, and individual cells (column) annotation for i) cell type final annotation as shown in (A), ii) sorting phenotype, iii) time point after MCMV infection, iv) belonging to R-phenograph clusters, and v) belonging to Seurat clusters. Six gene groups are shown: (1) correspond to gene specifically expressed at high levels in pDC, (2) corresponds to genes with shared selective expression in pDC and pDC-like cells, (3) are cDC1-specific genes, (4) are genes previously reported to be expressed at higher levels in pDC-like cells over pDC, (5) are cDC2-specific genes, and (6) are genes expressed selectively at higher levels in tDC and cDC2 or cDC1 or pDC-like cells. (D) Violin plots showing the expression of the tdT fluorescent protein across the DC types identified in (A). (E) Violin plots showing the showing mRNA expression levels of selected genes across DC types.

### pDC-Tomato mice allow mapping pDC localization in lymphoid and non-lymphoid organs

Since flow cytometry on cell suspensions from various organs and scRNA-seq analyses on splenocytes confirmed highly specific and penetrant expression of tdT in pDC from pDC-Tomato mice, we next harnessed this novel and unique pDC-reporter mouse model to determine the micro-anatomical localization of pDC in various organs. In the spleen, a relatively high density of tdT^+^ cells was observed in the T cell zone (TCZ) of the white pulp (WP) marked by a high density of CD3 staining (Figures 5A). Scattered tdT^+^ cells were also detectable in red pulp (RP) areas delimited by CD169^+^ metallophilic marginal zone (MZ) macrophages and encompassing a high density of F4/80^+^ cells, most of which are known to be Red Pulp Macrophages (Figure 5B). As expected, the overwhelming majority of tdT^+^ splenocytes expressed BST2, both in the TCZ and RP (Figure 5A-B). However, whereas most BST2^+^ cells in the TCZ were tdT^+^ (Figure 5A), this was not the case in the RP (Figure 5B), consistent with known expression of BST2 on other steady state splenocytes, including plasma cells, subsets of macrophages, pDC-like cells and tDC^7, 9, 10^. We then quantified the density of pDC (tdT^+^ cells) in the whole spleen (Figure 5C-D), as well as in its different micro-anatomical areas, the RP, the marginal zone (MZ) that delineates the white pulp (WP), and within the WP, the TCZ and the B cell zone (BCZ) (Figures 5C, E-F). The overall density of pDC in the whole spleen was around 400 cell/mm^2^ (Figure 5D). Around 40% and 30% of splenic pDC were located in the TCZ and RP, respectively, with only smaller fractions of pDC in the BCZ and MZ (Figure 5E). Remarkably, however, the density of pDC was much higher in the TCZ (∼1500 cells/mm^2^) than in all other splenic compartments that harboured similar values, at least 3.5 fold lower (≤400 cells/mm^2^) (Figure 5C, F). Consistent with their distribution in the spleen, in lymph nodes the pDC were highly frequent in paracortical CD3^+^ T cell areas (Figure S6A). Based on the distribution of cell surface markers thought to be selectively expressed by pDC, such as BST2, previous publications reported that in the small intestine pDC were present both in the Peyer’s patches^28^ and in the lamina propria (LP)^29^. However, it has been recently shown that other cell types express BST2 in this micro-anatomical compartment, in particular lysozyme M-expressing macrophages and DC^30, 31^. Hence, BST2 expression cannot be used reliably to determine the micro-anatomical location of pDC in the gut. Therefore, we analysed tissue sections of the small intestine from pDC-Tomato mice to re-examine the micro-anatomical location of pDC in this organ. tdT^+^ cells were detectable in the LP, preferentially in areas proximal to the EpCAM^+^ epithelial cell layer (Figure S6B). Only very few tdT^+^ cells were observed in Peyer’s patches (Figure S6C). Differently from the small intestine, pDC were not detectable in the LP of the large intestine (Figure S6D). These results indicate that, at steady state, in the gut, *bona fide* pDC are detectable exclusively in the LP of the villi of the small intestine; they are absent from the large intestine and from the organized lymphoid structures of the gut including Peyer’s patches.

**Figure 5.**
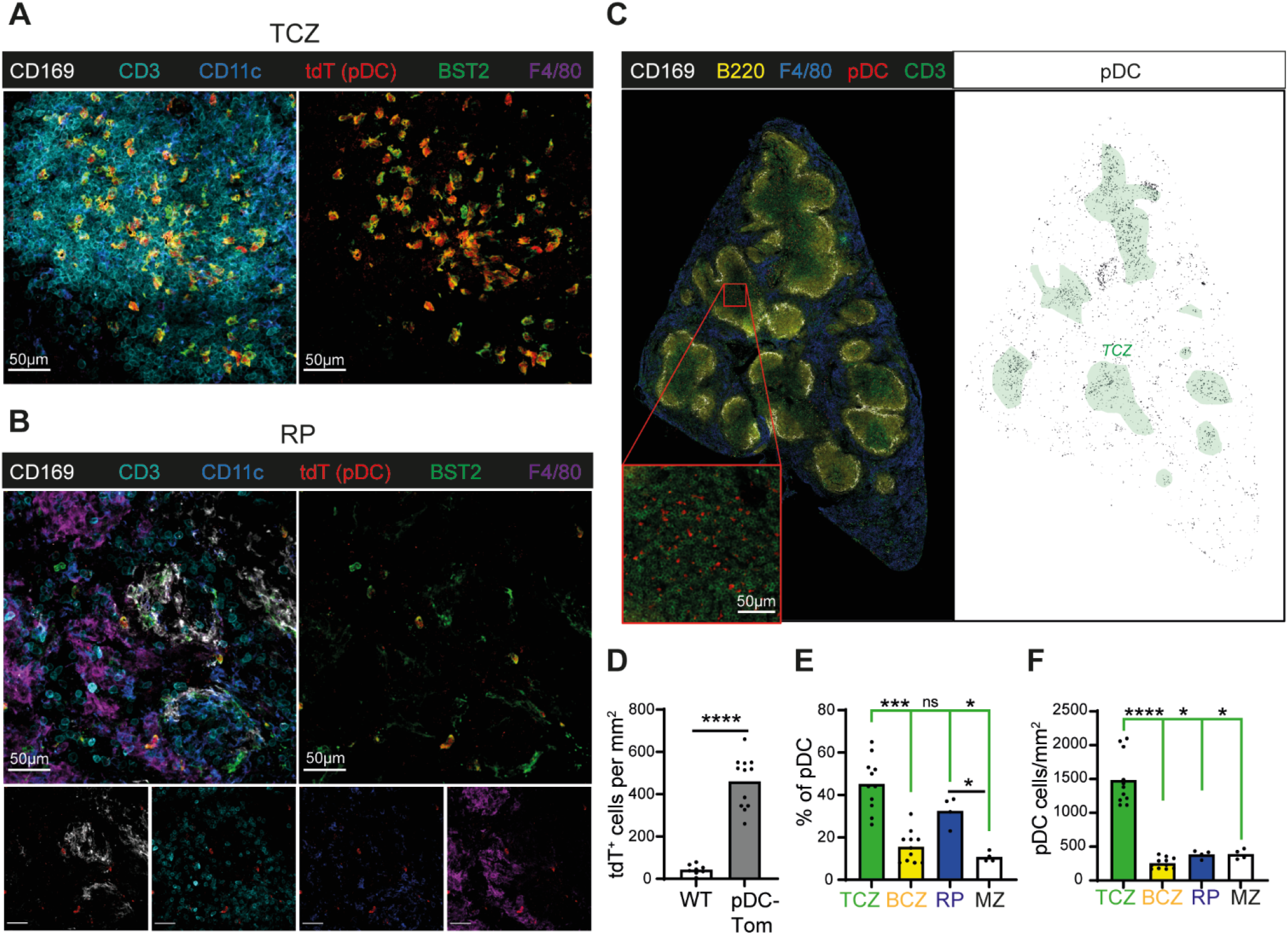
pDC-Tomato mice allow studying pDC micro-anatomical location in the spleen. 20 μm spleen cryosections from steady state pDC-Tomato mice were were stained with anti-tdT (red), anti-CD169 (white), anti-CD3 (cyan) and anti-F4/80 (violet or blue) antibodies, combined with anti-CD11c (blue) and anti-BST-2 (green) antibodies (A-B) or with an anti-B220 (yellow) antibody (C-F). (A-C) Representative images of 4 sections from 2 animals are shown, for the T-cell Zone (TCZ) region (A), the Red Pulp (RP) region (B), and the whole section (C) with the left panel corresponding to the raw signal and the right panel to the pDC mask used for quantification (black) relative to the location of the TCZ (delimited by the green lines, based on anti-CD3 staining). (D) The number of tdT^+^ cells per square millimeter of the whole spleen sections was quantified in pDC-Tomato mice, in comparison to WT animals as background controls. The data shown are from 8 whole sections from 2 WT mice and 9 whole sections from 3 pDC-Tomato mice. (E-F) Micro-anatomical distribution of pDC across the different area of the spleen, namely the TCZ (CD3-rich region), B-cell Zone (BCZ, B200-rich region), RP (F4/80-rich area) and Marginal Zone (MZ, defined as the space between the CD169 and F4/80 stainings). (E) Fraction of the pDC population present in each micro-anatomical area (calculated as the ratio between the pDC counts in the area to the total pDC counts in the whole section). (F) pDC density (cells/mm^2^) in each micro-anatomical area. The data shown are from 11 sections from 3 mice for TCZ and BCZ, and from 4 sections from 3 mice for MZ and RP. The height of the colored boxes shown mean expression across individual values for each micro-anatomical area. 1-way ANOVA was used for the statistical analysis, *P < 0.05, **P < 0.01, ***P < 0.001, ****P < 0.0001.

### SCRIPT mice allow comparing the spatiotemporal repositioning of the pDC producing or not IFN-I in the spleen during MCMV infection

By combining the use of *Ifnb1*^EYFP^ reporter mice^32^ with scRNA-seq, flow cytometry and confocal microscopy^24^, we recently determined the activation trajectory of splenic IFN-I-producing pDC *in vivo* during MCMV infection. In particular, we showed their clustering in the MZ near to infected cells at the time of peak IFN-I production, followed by their migration to the TCZ after termination of their IFN-I production and acquisition of transcriptomic, phenotypic and functional features of mature DCs. However, the lack of adequate markers prevented us from assessing the spatiotemporal repositioning of the pDC that were not producing IFN-I. To answer this question, we bred pDC-Tomato mice with *Ifnb1*^EYFP^ reporter animals, hence generating ***S****iglecH*^i^**^Cr^**^e^;***I****fnb1*^EYFP^;***P****acsin1^LSL-^****^t^****^dT^* mice, hereinafter called SCRIPT mice (Figure 6A). In SCRIPT mice, we can thus unambiguously identify both the pDC that produce IFN-I (tdT^+^ YFP^+^) and those that do not (tdT^+^ YFP^neg^), for the first time to our knowledge (Figures 6A-B). By flow cytometry, both tdT^+^ YFP^neg^ and tdT^+^ YFP^+^ cells were detectable in splenocyte suspensions 36 and 48 hours after MCMV infection, with tdT^+^ YFP^+^ representing 20-25 % of the total splenic pDC (tdT^+^) population (Figure 6B). Both populations expressed comparable high levels of Ly6D and BST2, consistent with their pDC identity (Figure 6B). We also detected rare tdT^neg^ YFP^+^ Ly6D^neg^ BST2^low^ cells. These data are consistent with our previous work^5, 17, 24, 33^, confirming that pDC are the main source of IFN-I during MCMV infection, with other cell types only marginally contributing to this production. Next, we examined the localization of both tdT^+^ YFP^neg^ and tdT^+^ YFP^+^ pDC in spleens from MCMV-infected mice, by using confocal microscopy (Figures 6C-F). At 36h and 40h after infection, both subsets of pDC formed clusters in the MZ (Figure 6C-D) leading to an increase of the proportion of splenic pDC at this micro-anatomical site (Figure 6E). Although pDC were still detectable in the TCZ, their local density was reduced as compared to uninfected mice (Figure 6C, E). At 48h after infection, we detected large aggregates of tdT^+^ YFP^neg^ cells in the MZ, whereas tdT^+^ YFP^+^ cells were most prominently detected in the TCZ (Figure 6C-D). Hence, we observed an opposite trend in the spatiotemporal repositioning of the pDC producing or not IFN-I within the spleen during MCMV infection: whereas the majority of IFN-I-producing (YFP^+^) pDC were located in the MZ at 36h but then in the TCZ at 48h, the reverse distribution was observed for the pDC that did not produce IFN-I (YFP^neg^) (Figures 6F). This analysis showed for the first time to our knowledge that, at the peak of IFN production, the high recruitment of pDC to the MZ occurs independently of their ability to produce IFN, whereas, at later time points, only the pDC that produced IFN-I are licensed to migrate to the TCZ while the other pDC are retained in the MZ. Finally, we sorted the different DC from the spleens of infected mice and examined their morphology using confocal microscopy, also comparing YFP^+^ versus YFP^neg^ pDC from mice that had been infected for 60h (Figure 6G-H and Figure S7). This analysis demonstrated a clear and selective change in the morphology of YFP^+^ pDC, a significant fraction of which acquired an irregular morphology with several pseudopods or dendrites (Figure 6G), translating into significantly lower circularity indices as compared not only to pDC from uninfected animals but also to the YFP^neg^ pDC isolated from the same infected mice (Figure 6H). Hence, this quantitative and unbiased confocal microscopy analysis demonstrated the acquisition of a classical dendritic cell morphology selectively by the pDC that had produced IFN-I during MCMV infection, consistent with our previously reported demonstration of their transcriptomic and functional convergence towards DC^24^.

**Figure 6.**
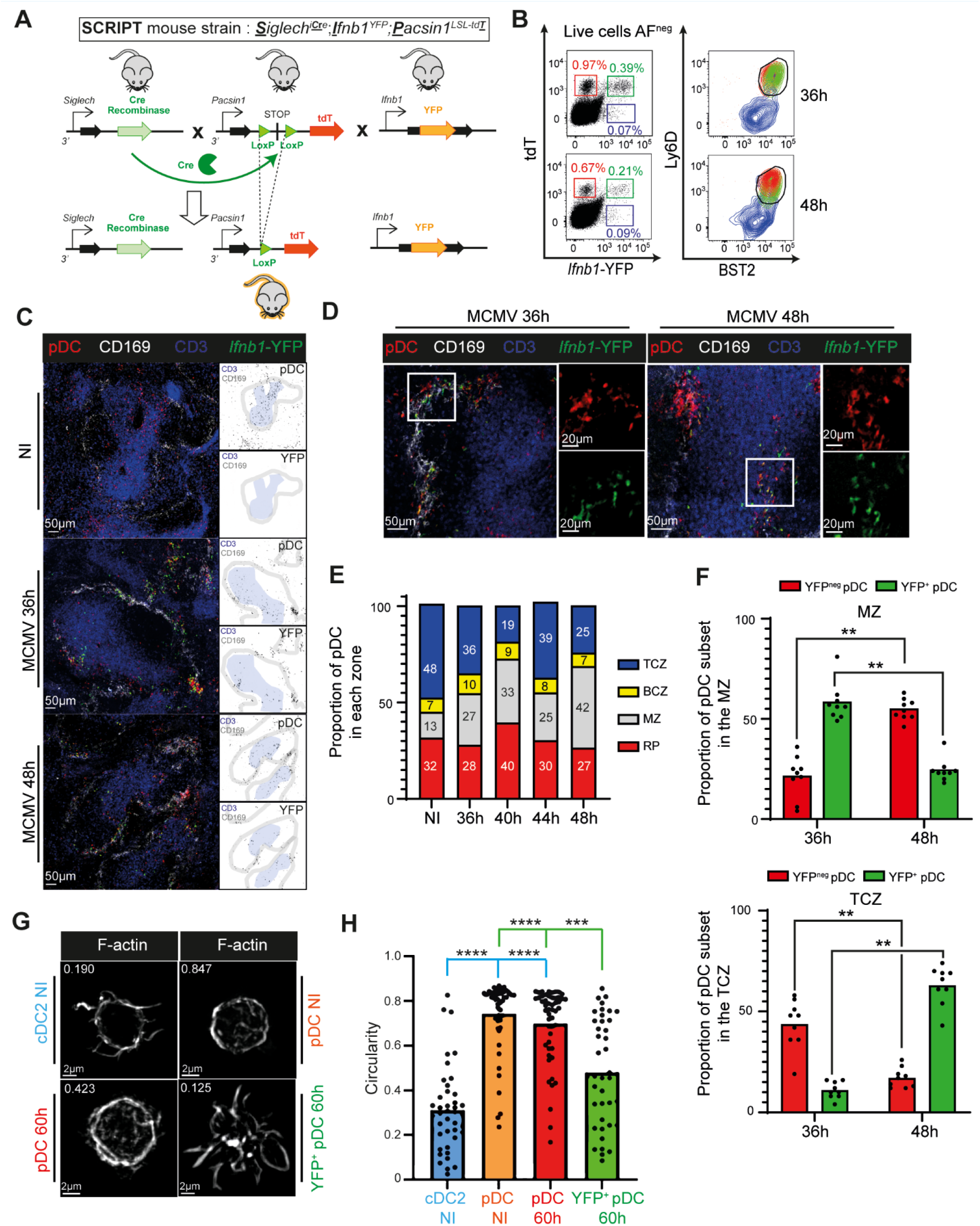
Diverging intra-splenic migration patterns and morphological changes between IFN-producing versus non-producing pDC during MCMV infection. (A) Scheme representing the strategy followed to generate SCRIPT mice. (B) Phenotypic characterization of *Ifnb1*-expressing cells in the spleen of SCRIPT mice at 36h and 48h after MCMV infection, by flow cytometry. Within live, non-autofluorescent (AF^neg^), cells, pDC were identified as producing IFN-I (YFP^+^ tdT^+^, green boxes and contour plot) or not (YFP^neg^ tdT^+^, red boxes and contour plots), and other IFN-I producing cells were identified as tdT^neg^ YFP^+^ (violet boxes and contour plots); their expression of Ly6D and BST2 was then examined (right contour plots). (C-F) 20 μm spleen cryosections from pDC-Tomato mice infected by MCMV for 36h or 48h were stained with anti-CD169 (white), anti-CD3 (blue), anti-tdT (red), and anti-YFP (green) antibodies. (C) The scheme on the right of each photograph shows the masks used for quantification, with pDC shown in black, the TCZ delimited as the blue areas, and the MZ as the gray area. (D) Representative images of a pDC cluster in the MZ at 36h post-infection (left) and of the selective migration of the YFP^+^ pDC in the TCZ at 48h post-infection (right). (E) Micro-anatomical distribution of pDC across the different area of the spleen, during the course of MCMV infection. The data shown are from 6 animals for uninfected mice, 9 animals for 36h, 7 for 40h, 8 for 44h and 9 for 48h, with one whole spleen section analysed per mouse. (F) Fraction of the YFP^+^ versus YFP^neg^ pDC populations present in the MZ (left) or TCZ (right) at 36h versus 48h post-infection. The data shown are from 9 mice for each time point, from the same animals as shown in (E). The height of the colored boxes shown mean expression across individual values for each micro-anatomical area. A Wilcoxon t test was used for the statistical analysis, \**P <* 0.05, \*\**P <* 0.01. (G-H) Quantitative and unbiased assessment of the cellular morphology of YFP^+^ versus YFP^neg^ pDC isolated from 60h MCMV-infected SCRIPT mice, as compared to pDC and cDC2 from uninfected mice. (G) One representative confocal microscopy image of each DC type is shown. (H) The distribution of the circularity indices for individual cells across DC types is shown as dots, with the overlaid color bars showing the mean circularity indices of each DC type. The data shown are from 2 independent experiments each performed with one mouse, with 42 to 53 individual cells analyzed for each DC type. The data are shown as mean ± SEM. Kruskal-Wallis was used for the statistical analysis, *P<0.05, **P<0.01, ***P<0.001, ****P<0.0001.

### The recruitment of pDC to the vicinity of infected cells in the marginal zone of the spleen occurs early after MCMV infection

Due to the lack of specific enough markers for pDC before their peak production of IFN-I at 36h after virus inoculation, it has previously not been possible to determine the early kinetics of their micro-anatomical redistribution in the spleen during the course of MCMV infection. The generation and validation of the pDC-Tomato mice allowed us to address this issue for the first time to our knowledge (Figure 7). The recruitment of pDC and their clustering in the MZ of the spleen were detectable as early as 12h after infection (Figure 7A-B), with a clear increase in the proportion of pDC in the MZ close to statistical significance and to the plateau values observed between 18h and 48h (Figures 6E and 7C). Cells expressing the viral immediate early gene 1 (IE1) and thus replicating MCMV were also already detectable at 12h after infection, mainly in the MZ (Figure 7D-F). The pDC clusters observed in the spleen of infected mice were found in close proximity with MCMV-infected cells at all the time points examined, including already at 12h (Figure 7D), consistent with our previous findings at later time points^17^. Thus, these analyses demonstrate that the recruitment of pDC to the vicinity of infected cells in the marginal zone of the spleen occurs early during MCMV infection, already at 12h after virus inoculation, thus 24h before their peak IFN-I production.

**Figure 7.**
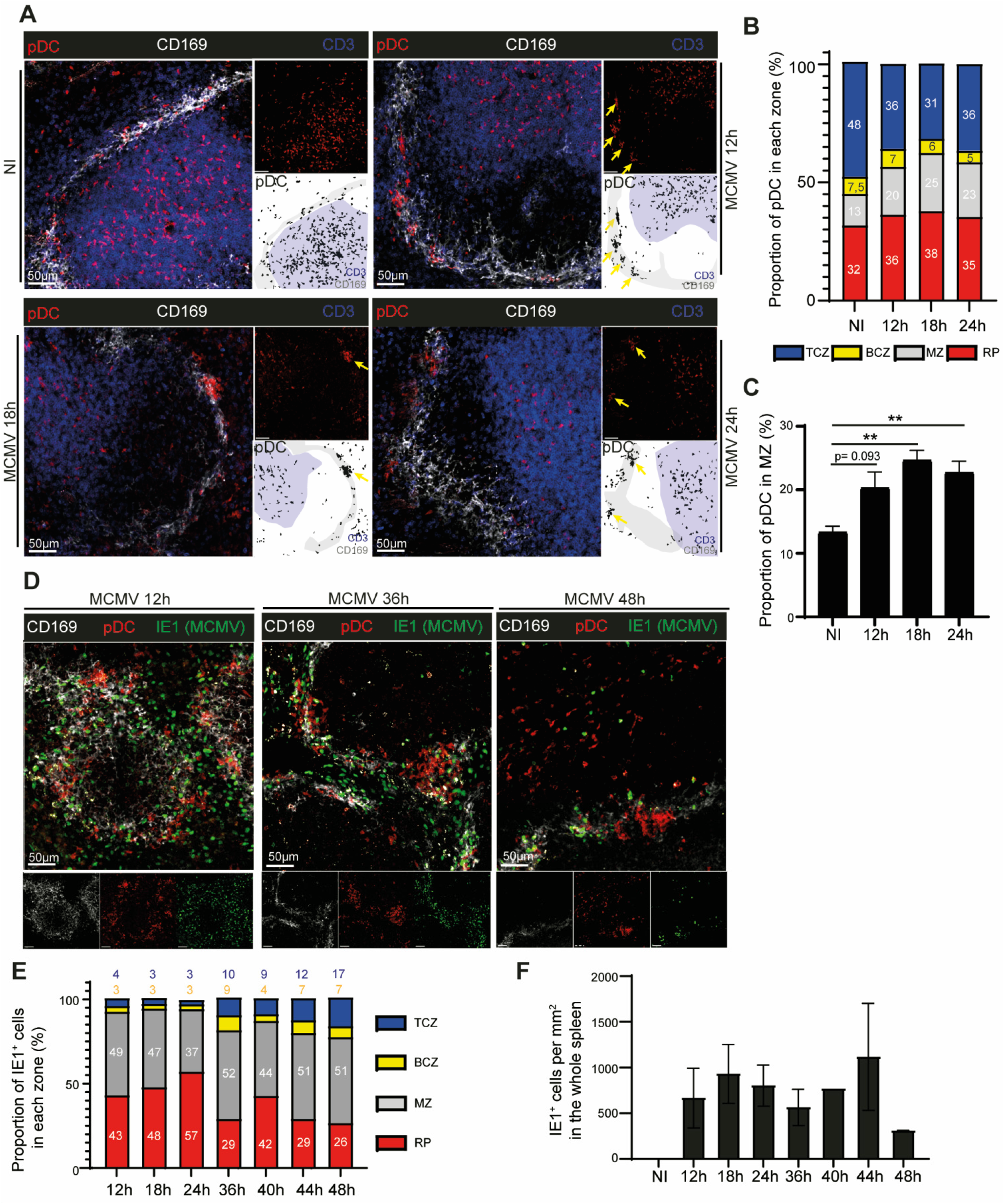
Early during MCMV infection pDC are recruited at the MZ where they contact infected cells. 20μm spleen cryosections from pDC-Tomato mice were stained with anti-tdT (red), anti-CD169 (white) and anti-CD3 (blue) antibodies (A-C) or with anti-tdT (red), anti-CD169 (white) and anti-IE1 (green) antibodies (D-F). (A) Representative images for NI or 12h, 18h and 24h MCMV infection conditions. (B) Micro-anatomical distribution of pDC across the different areas of the spleen, during the course of MCMV infection. The data shown are from 6 animals for uninfected mice, 6 animals for 12h, 9 for 18h, and 11 for 24h, with one whole spleen section analysed per mouse. (C) The proportion of pDC in the MZ was quantified. The data are shown as mean±SEM. 1-way ANOVA was used for the statistical analysis, *P<0.05, **P<0.01, ***P<0.001, ****P<0.0001. (D) Representative images for NI or 12h, 36h and 48h MCMV infection conditions. (E) Micro-anatomical distribution of IE1^+^ cells across the different areas of the spleen, during the course of MCMV infection. The data shown are from 2 animals for uninfected mice, 2 animals for 12h, 2 for 36h, and 2 for 36h, with one whole spleen section analysed per mouse. (F) The number of IE1^+^ cells per mm^2^ was quantified in the whole spleen section. The data are shown as mean±SEM.

### scRNA-seq confirms the unique capacity of pDC for high IFN-I/III expression during MCMV infection and shows divergent activation patterns for pDC-like cells and tDC

Finally, we analysed our FB5P-seq data to compare the responses of the five DC types to MCMV infection *in vivo*. Four Seurat clusters were identified for pDC, corresponding to 4 activation states (Figure 8A): quiescent pDC (mostly from uninfected animals), intermediate pDC harbouring induction of interferon-stimulated genes (ISG) but lacking expression of cytokines genes, activated pDC expressing moderate levels of cytokine genes, and IFN-I-producing pDC expressing high levels of all of the genes encoding IFN-I. tDC, cDC1 and cDC2 were each split into three Seurat clusters, corresponding to quiescent, intermediate and activated states. Only two activation states were observed for pDC-like cells, quiescent and activated. Importantly, tdT expression remained high selectively in pDC, irrespective of their activation states (Figure 8B), despite the decreased SiglecH expression occurring in IFN-I-producing pDC (Figure S8A)^5^. Activation increased autofluorescence in other DC types but did not induce their expression of tdT (Figure 8B). pDC also remained characterized by their uniquely high levels of Ly6D and CCR9 expression as compared to the other DC types, across activation states. In contrast, BST2 expression increased on all activated DC types (Figure 8B), as reported previously^5, 34, 35^, leading to a strong overlap of BST2 expression levels between activated pDC and pDC-like cells. Activation decreased CX_3_CR1 expression in tDC (Figure S8), which, in conjunction with the increased BST2 expression (Figure 8B), made their phenotypic discrimination from pDC-like cells more difficult, although CD11c and GFP expression remained higher on tDC than on pDC-like cells (Figure S8A). Although XCR1 expression on cDC1 and CD11b on cDC2 decreased with activation, they remained clearly detectable, still allowing phenotypic identification of these DC types (Figure S8A).

**Figure 8.**
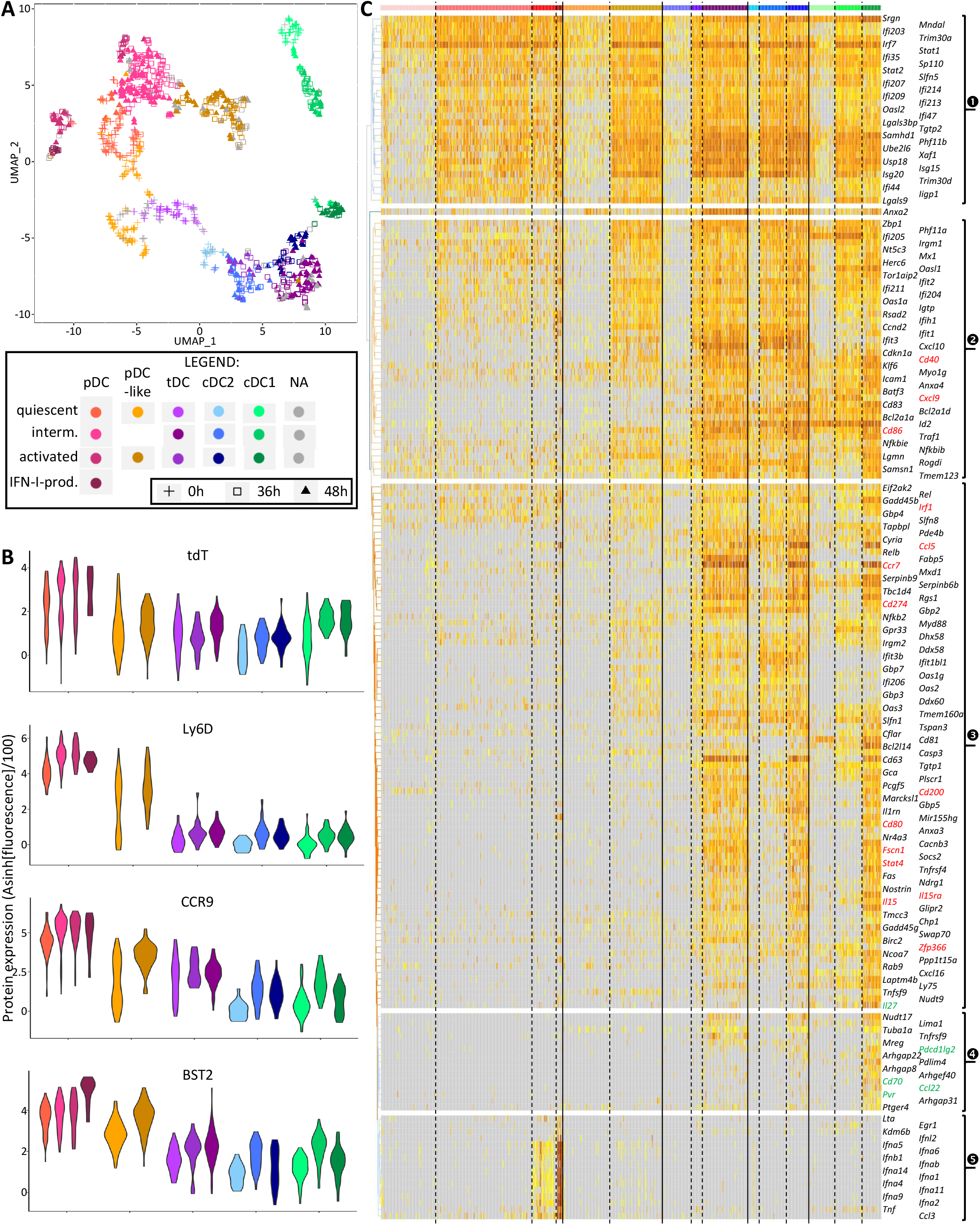
scRNA-seq confirms the unique capacity of pDC for high IFN-I/III expression during infection and shows divergent activation patterns for pDC-like cells and tDC. (A) Projection of assigned DC type and activation states (color code) onto the UMAP space based on FB5P-seq gene expression. DC type assignment is the same as in Figure 4A. Activation states were assigned based on mining of the marker genes of Seurat clusters. (B) (B) Violin plots showing the expression of selected phenotypic markers across DC types and activation states. (C) Heatmap showing mRNA expression levels of selected genes (rows) across all 951 individual cells (columns), with hierarchical clustering of genes using Euclidian distance, and ordering of individual cells (column) according to their assignment into cell types and activation states using the same color code (top) as in panel (A). The color scale for gene expression levels is the same as in Figure 4C.

In the UMAP space, quiescent tDC and pDC-like cells were close to one another, between pDC and cDC2 (Figure 8A), as observed in previous reports in UMAP spaces drawn from flow cytometry or bulk RNA-seq data^9^. Strikingly, however, upon activation they moved towards opposite directions, with activated pDC-like cells close to intermediate pDC, but intermediate and activated tDC close to intermediate and activated cDC2 (Figure 8A). Hence, this analysis suggested divergent activation patterns for pDC-like cells and tDC *in vivo* during MCMV infection. Indeed, whereas all DC types induced expression of many ISG (Figure 8C, cluster 1), only tDC, cDC2 and cDC1 induced high levels of genes associated to DC maturation and linked to their migration to the T cell zones of lymphoid organs and to cognate interactions with T cells, encompassing *Fscn1*, *Ccr7*, *Il15*, *Il15ra*, *Cd80*, *Cd200*, *Cd274* (Figure 8C, cluster 3). Another set of ISG and of genes associated to DC interactions with T cells was induced to gradually increasing levels from intermediate to activated to IFN-I-producing pDC (Figure 8C, cluster 2), consistent with our recent characterization of the activation trajectory of these cells^24^. These genes were induced to similar levels in IFN-I-producing pDC and in activated pDC-like cells, but to even higher levels in activated tDC, cDC2 and cDC1. Only few genes were found to be differentially expressed between tDC and cDC2: 16 at steady state, 1 at the intermediate activation state, and 11 for activated cells. Whereas only 84 genes were differentially expressed between tDC and pDC-like cells at steady state, this number increased to 527 for activated cells. In contrast, the number of genes differentially expressed between pDC and pDC-like remained stable over activation, 113 at steady state versus 138 in activated cells. Hence, this analysis confirmed the divergent activation patterns for pDC-like cells and tDC suggested by the UMAP, with pDC-like cells behaving closer to pDC, although not producing IFN-I (Figure 8C, cluster 5), whereas tDC converging further towards cDC2.

Expression of the *Ly6c2* gene remained much higher in pDC-like cells than in tDC (Figure S8B), suggesting that the distinction between pDC-like cells and tDC based on their reciprocal expression levels of CD11c (Figure S8A) and Ly6C as observed in steady state^9^ could hold under activation conditions. Genes encoding MHC-II molecules were consistently higher in tDC than in pDC-like cells, reaching the same levels as observed in cDC2, as illustrated for *H2-DMb2* and *H2-Eb1* (Figure S8B). The genes *Tmem176a* and *Tmem176b*, encoding cell surface markers, were consistently and significantly higher in tDC than in pDC-like cells, pDC and cDC1 across activation conditions and also tended to be lower in cDC2 than tDC (Figure S8B). Hence, we here identified Tmem176a and Tmem176b as candidate markers specific for tDC. Expression of *Cd8a* remained lower in cDC2 than in other DC types across activation conditions (Figure S8B). The gene *Ms4a4c* was higher in activated pDC-like cells, intermediate and activated tDC and cDC2 than in pDC and cDC1 (Figure S8B). One gene, *Apod*, was exclusively expressed to high levels in pDC-like cells above all other DC types across activation conditions (Figure S8B).

Finally, beyond confirming the unique ability of pDC to produce IFN-I/III during MCMV infection^5, 17, 33^, our analysis also unravelled a specific response of cDC1 that selectively induced high expression of several genes including positive or negative co-stimulation molecules and cytokines involved in the cross-talk with NK cells and T lymphocytes, encompassing *Cd70*, *Pdcd1lg2*, *Pvr*, *Ccl22* (Figure 8C, cluster 4), *Il27* (Figure 8C, cluster 3) *Il18* and *Il12b* (Figure S8B). This is consistent with a division of labor between DC types, with cDC1 playing a unique and non-redundant role in promoting NK cell antiviral activity and accelerating CD8^+^ T cell responses during MCMV infection^36–39^.

## Discussion

In a review on how genetic models of DC development enable correlating ontogeny with function, Pr. Kenneth Murphy, an international leader in this research field, stated that “pDC function is still a black box” because “a model of pDC-restricted functional or developmental impairment [remains to be] developed”^6^. Indeed, the lack of mutant mouse models enabling to specifically target pDC has hampered our understanding of their physiological functions, their choreography *in vivo* and their molecular regulation. This is all the more the case since, when classical markers are used for identification, pDC can be contaminated with other cell types reported to be much more powerful than resting pDC for T cell activation^7–9^, including pDC-like cells and tDC. This recent discovery has put into questions some of the conclusions previously drawn on pDC, because these cell types can also be affected in the experimental settings previously used to deplete pDC. The injection of anti-GR1 or –Ly6C or –BST2 antibodies should especially affect pDC-like cells, in addition to subsets of neutrophils, monocytes or macrophages. The use of the CD11c-Cre;Tcf4-fl/fl mouse model also leads to tDC depletion^9^. This would also be expected in SiglecH-DTR mice where other cell type are already known to be affected, including DC and subsets of macrophages^14^. For the same reason, and because pDC downregulate SiglecH upon activation, the use of Siglech-GFP reporter mice to track pDC *in vivo* is inadequate^14^. Although GFP expression is high and penetrant in pDC from the DPE^GFP^ mice^40, 41^, this model is inadequate to track pDC *in vivo,* since T and NK T cells also strongly express GFP that is under the control of the distal and proximal CD4 enhancers and CD4 promoter^42, 43^. Moreover, other myeloid cells express the *Cd4* gene including subsets of cDC2 and of macrophages^30^. Thus, novel tools are required to specifically identify, characterize and manipulate pDC *in vivo*, including to discriminate them from tDC that strongly infiltrate inflamed tissues^9, 44^. However, this aim has been difficult to achieve because no single gene has been found to be expressed specifically in pDC. Hence, to overcome this bottleneck, we devised an intersectional genetic strategy based on the unique co-expression of *Siglech* and *Pacsin1* in pDC^15^. Indeed, whereas Siglech is also expressed in pDC-like cells, DC precursors and subsets of macrophages, this is not the case of *Pacsin1* whose expression is otherwise confined to neurons. Hence; we report here the generation and characterization of pDC-reporter mice as *Siglech^iCre^*;*Pacsin1^LSL-tdT^* animals.

We show high specificity and penetrance of tdT expression in pDC, by a combination of methods including flow cytometry, confocal microscopy, and scRNA-seq, in different organs and both at steady state and during the inflammation induced by MCMV infection *in vivo*. Using this new mouse model, for the first time to our knowledge we could study the choreography of the relocation of all pDC in the spleen during MCMV infection. We show that many pDC clustered in the marginal zone in the vicinity of infected cells early after infection, but that their fate diverged later on depending on their activation state. The pDC failing to produce IFN-I remained in the marginal zone, whereas the pDC that had produced IFN-I migrated to the T cell zone. This shows that all pDC are attracted to the micro-anatomical sites of viral replication early after infection. Thus, restricted access to contact with infected cells is likely insufficient to explain how the majority of pDC fail to produce IFN-I. This raises the hypothesis that the pDC that produced IFN-I the first might have repressed this function in the other pDC present in the same cell clusters, through some yet undefined quorum sensing mechanism, to prevent excessive inflammation and the ensuing immunopathology. Further studies using pDC-reporter mice and innovative tools for pDC-specific genetic manipulations should help better understand this tight regulation.

Crossing our pDC-reporter mouse model with the *Zbtb46^GFP^* reporter mice enabled us to generate the first side-by-side transcriptomic comparison of pDC, pDC-like cells and tDC, through scRNA-seq, both at steady state and during MCMV infection, also including cDC1 and cDC2 as key reference DC types. Our results confirmed that only pDC produce high levels of all IFN-I *in vivo* during MCMV infection. We determined for the first time to our knowledge that pDC and pDC-like cells underwent divergent activation patterns, with only tDC showing features of classical DC maturation akin to that occurring in cDC2 and cDC1. We report the identification of gene modules induced during activation selectively in a given DC type, or in specific combinations of DC types. This dataset should be a valuable resource to mine in the coming years to infer from their specific gene expression programs hypotheses on the functional specialization of pDC-like cells and tDC, as compared to pDC, cDC1 and cDC2, at steady state and during a viral infection *in vivo*, and on their molecular regulation. We identified Tmem176a and Tmem176b as novel putative cell surface markers specific to tDC, which should help studying these cells in a more specific manner to understand their functions. Interestingly, these molecules have been reported to promote MHC-II antigen presentation to naïve CD4 T cells^45^ and MHC-I antigen cross-presentation to CD8 T cells, in a manner that may contribute to immunosuppression in tumors^46^ or to tolerance to allografts^47, 48^. Interestingly, Tmem176a/b were shown to strongly co-localize with HLA-DM in the late endolysosomal system, and, here, we show that tDC and cDC2 express higher levels of both *Tmem176a/b* and *H2.DMb2* than other DC types. This suggests the hypothesis of a possible functional specialization of tDC in the induction of CD4 T cell tolerance, and raises the question of the division of labor between tDC and cDC2. Based on the specific gene modules that we have identified here, these questions might be addressed in future studies, once mutant mouse models have been designed to specifically target tDC or cDC2.

Finally, our work opens the exciting perspective to now generate new mutant mouse models for specific and penetrant constitutive ablation of pDC, as *Siglech^iCre^*;*Pacsin1^LSL-Dta^* mice. It will also be exciting to try harnessing this intersectional genetic strategy to engineer a mouse model for binary Cre-mediated genetic editing specifically in pDC^49^. These mouse models should then be key to allow us better understanding the physiological functions of pDC in future studies.

## Materials and Methods

### Mice

All animal experiments were performed in accordance with national and international laws for laboratory animal welfare and experimentation (EEC Council Directive 2010/63/EU, September 2010). Protocols were approved by the Marseille Ethical Committee for Animal Experimentation (registered by the Comité National de Réflexion Ethique sur l’Expérimentation Animale under no. 14; APAFIS#1212-2015072117438525 v5 and APAFIS#21626-2019072606014177 v4). C57BL/6 mice were purchased from Janvier Labs, France. All other mouse strains were bred at the Centre d’ImmunoPhénomique (CIPHE) or the Centre d’Immunologie de Marseille-Luminy (CIML), Marseille, France, under specific pathogen free-conditions and in accordance with animal care and use regulations. *Siglech^iCre^* mice (*B6-Siglech^tm1(iCre)Ciphe^*)^17^ and *Pacsin1^LoxP-STOP-LoxP-tdTomato(LSL-tdT)^* (*B6-Pacsin1^tm1(tdT)Ciphe^*) mice were generated by CIPHE (Centre d’Immunophénomique), Marseille, France. *Siglech^iCre^;Pacsin1^LSL-tdT^* mice (pDC-Tomato) were generated by crossing *Siglech^iCre^* mice with *Pacsin1^LSL-tdT^* mice, and then maintained and used at a double homozygous state. *Siglech^iCre^;Ifnb1^EYP^*;*Pacsin1^LSL-tdT^* (SCRIPT) mice were generated by crossing pDC-tomato mice with *Ifnb1^Eyfp^* mice (*B6.129-Ifnb1^tm1Lk^*^y^)^32^, and then maintained and used at a triple homozygous state. *Zbtb46^GFP^;Siglech^iCre^;Pacsin1^LSL-tdT^* (ZeST) mice were generated by crossing *Zbtb46^GFP^* mice (*B6.129S6(C)-Zbtb46^tm1.1Kmm/J^*)^23^ with pDC-Tomato mice and were used at a heterozygous state. All animals used were sex- and age-matched (used between 8-16 weeks of age).

### Virus and viral infection

Virus stocks were prepared from salivary gland extracts of 3-weeks old MCMV-infected BALB/c mice. All mice used in the experiments were infected i.p. with 10^5^ pfu Smith MCMV and sacrificed at indicated time points.

### Cell preparation for flow cytometry analysis or cell sorting

Spleens or lymph nodes were harvested and submitted to enzymatic digestion for 25 minutes at 37°C with Collagenase IV (Wortington biochemicals) and DNAse I (Roche Diagnostics). Organs were then mechanically digested and passed over 100µm cell strainers (Corning). Red blood cells were then lysed by using RBC lysis buffer (Life Technologies) for spleen and bone marrow cell preparation. Livers were harvested, minced and submitted to enzymatic digestion, as for the spleen. Liver pieces were then crushed and cell suspension obtained was washed 2 times with PBS 1x, before performing a 80/40 Percoll gradient. Cells isolated from the middle ring of the gradient were washed once with PBS 1x, then used for flow cytometry. Small intestines were harvested, opened longitudinally, then cut into 1mm pieces. Pieces were washed entensively with PBS 1x, then incubated 3 times at 37°C upon shaking (200 rpm) with PBS 1X containing 2% Fetal Calf Serum (FCS) and 5mM ethylenediamine tetraacetic acid (EDTA). At the end of each incubation, supernatants were collected and centrifuged. Pelleted cells, mainly IntraEpithelial Cell (IEL), from the three incubations were pooled together and submitted to a 67/44 Percoll gradient. Cells isolated from the middle ring of the gradient were washed once with PBS 1x, then used for flow cytometry.

### Flow cytometry analysis

Extracellular staining was performed in PBS 1X supplemented with 2 mM EDTA (SigmaAldrich) and 1% FCS. All extracellular stainings were performed for at least 30 min at 4°C. Dead cell staining (LIVE/DEAD™ Fixable Aqua Dead Cell Stain or Blue Dead Cell Stain, Life Technologies) was performed in PBS 1X according to the manufacturer’s recommendations. Samples were acquired with a FACS Fortessa X20 (BD Biosciences) or sorted with FACS ARIA III (BD Biosciences). All data were analyzed with the Flowjo software. For the unsupervised analysis, tSNE plots were generated with the dedicated plugin in the software. Dimensional reduction was performed on Singlets/Non Autofluorescent/Live Dead^neg^/Lineage^neg^/CD19^neg^/CD11c^+^ and/or SiglecH^+^ cells. Dimension reduction was calculated with the following markers: B220, BST2, CD11b, CD11c, CCR9, CX_3_CR1, Ly6D, SiglecH, tdT, XCR1 and *Zbtb46*-GFP. For the generation of unsupervised gating strategy, the plugin Hyperfinder was used. The calculation was performed on Singlets/Non Autofluorescent/Live Dead^neg^ cells. We defined the pDC population for the calculation as being Lineage^neg^/CD19^neg^/ CD11b^neg^/XCR1^neg^/CD11c^low^/BST2^high^/tdT^+^. The software calculated the best and fastest gating strategy with the following markers: B220, BST2, CD11b, CD11c, CD19, CCR9, CX_3_CR1, Lineage (CD3/Ly6G/NK1.1) Ly6D, SiglecH and XCR1.

### scRNA-seq data generation

For the generation of the scRNAseq data, we followed the FB5P previously published method^25^. Briefly, single cells were FACS-sorted into ice-cold 96-well PCR plates (Thermofisher) containing 2µl lysis mix per well. Immediately after cell sorting, each plate was covered with an adhesive film (Thermofisher), briefly spun down in a benchtop plate centrifuge, and frozen on dry ice. The reverse transcription reaction was performed with SuperScript II (Thermofisher) in presence of RnaseOUT (Thermofisher), DTT (Thermofisher), betaine (Sigma-Aldrich), MgCl2 (Sigma-Aldrich), and well-specific template switching oligonucleotide. For cDNA amplification, KAPA HiFi HotStart ReadyMix (Roche Diagnostics) was used with adapted primers. For library preparation, amplified cDNA from each well of a 96-well plate were pooled, purified with two rounds of 0.6X solid-phase reversible immobilization beads (AmpureXP, Beckman, or CleanNGS, Proteigene) and finally eluted in nuclease free water. After tagmentation and neutralization, tagmented cDNA was amplified with Nextera PCR Mastermix containing Nextera i5 primer (Illumina), and custom i7 primer mix. Libraries generated from multiple 96-well plates of single cells and carrying distinct i7 barcodes were pooled for sequencing on an Illumina NextSeq2000 platform, with High Output 75 cycles flow cells, targeting 5 × 10^5^ reads per cell in paired-end single-index mode with the following cycles: Read1 (Read1_SP, 67 cycles), Read i7 (i7_SP, 8 cycles), and Read2 (Read2_SP, 16 cycles).

### Bioinformatics analyses of scRNAseq data

FB5P sequencing data were processed to generate a single-cell UMI counts matrix as described^25^. The counts matrix was loaded to R (v.4.0.3), and Seurat (v3.2.0)^50^ was used for downstream analyses as described^24^. Gene expression is shown as log-normalized values and protein expression as inverse hyperbolic arcsine (asinh) of fluorescence intensity scaled values (asinh(fluorescence/100)). For dimensionality reduction, we performed uniform manifold approximations and projections (UMAP), using the RunUMAP function. The differentially expressed genes were determined using the FindMarkers function.

A first Seurat analysis was performed only on cells from uninfected mice (345 cells after QC, 2 of which were removed due to lack of index sorting data). Clusters were identified based on gene expression using Seurat (k-nearest neighbor=5; resolution=0.2) or based on phenotypic marker expression using Rphenograph^51^ (v0.99.1) (number of nearest neighbours k=20), taking into account the protein expression for GFP, B220, Ly6D, XCR1, CX_3_CR1, BST2, SiglecH, CCR9, CD11b and CD11c (but not for tdTomato). A single cell connectivity Map (sgCMap) analysis was performed on this dataset using cell type-specific signatures (tDC, cDC2, cDC1, pDC) established upon re-analysis with BubbleGUM GeneSign module^52^ of a published^26^, independent, bulk RNA-seq dataset (GEO accession number GSE76132), and a relative pDC_vs_pDClike signature retrieved from a previously published study^7^.

Integration of Seurat and Phenograph cluster information together with the cMap scores allowed identifying, and focus on, *bona fide* steady state pDC, pDC-like cells, tDC, cDC1 and cDC2 (205 total cells), from which we computed relative signatures based on all pairwise comparisons between cell types, using the FindMarkers function (default parameters), for consecutive single cell cMap analyses.

A second Seurat analysis was then performed on cells from both uninfected and MCMV-infected mice (1132 cells after QC). Clusters of contaminating cell types (macrophages, NK cells, and a small cluster of proliferative cells of mixed types) were identified by their top markers using the FindMarkers function (test.use = “bimod”) and removed (181 cells). A third Seurat analysis was performed on the remaining cells (951 cells).

Clusters were identified based on gene expression using Seurat (k-nearest neighbor=9; resolution=0.7) or based on phenotypic marker expression using Rphenograph (number of nearest neighbours k=50). Integration of Seurat and Phenograph clusters allowed to identify unambiguous cell types and activation states (851 total cells), which was corroborated upon performing a single cell cMap analysis using the relative signatures identified in the previous step. The genes differentially expressed between cell types at equivalent activation states or between activation states for a given cell type were then extracted, using the FindMarkers function (default parameters, threshold for adjusted p-value<0.05).

The heatmaps were plotted using the Gene-E program.

### Immunohistofluorescence, microscopy and image analysis

Organs were fixed with Antigen Fix (Diapath) for 2 hours for the small intestine, colon, lymph nodes or 4 hours for the spleen at 4°C, and then washed several times in PB (0,025 M NaH_2_PO_4_ and 0,1 M Na_2_HPO_4_). Organs were then immerged in a solution of 30% sucrose O/N at 4°C. Organs were then embedded in OCT (Sakura), snap frozen and stored at −80°C. 20μm cryosections were performed with a microtome (Leica 3050s Cryostat) at temperatures between −20°C to −22°C. For immunostainings, sections were blocked with PB 0.1% TritonX100 2% Bovine Serum Albumin (BSA) for 30 min at room temperature and then stained overnight at 4°C with primary antibodies diluted in PB 0.1% TritonX100 2% BSA. After several washings with PB, sections were then stained in PB 0.1% TritonX100 2% BSA with secondary antibodies 2 hours at 4°C. To analyze YFP signal, after the incubation with secondary antibodies, sections were washed and incubated with PB 0,1% TritonX100 2% BSA 5% Rabbit Serum for 30 min at room temperature. Sections were then stained with anti-GFP antibodies directly labeled with Alexa 488 for 2 hours at 4 °C. Finally, sections were washed with PB and mounted with a coverslip and Prolong Antifade Gold mounting medium (Life technologies). Whole sections were acquired by spectral confocal microscope (Zeiss LSM 880) with 20x or 40x objectives. Pictures were then analyzed with ImageJ, including through the development of several specific macros available on demand to the corresponding authors. The T cell zone was defined as a CD3-rich region within the white pulp. The B cell zone was defined as a B220-rich area within the white pulp. The red pulp was defined as an F4/80-rich region and the marginal zone was defined as the space between the CD169 and F4/80 stainings. For the calculation of pDC counts per mm^2^, the function “analyze particles” was used and a threshold for tdT intensity and the size (above 8μm^2^) was applied. For the quantification of the repartition of pDC upon infection, the intensity of tdT was calculated in the different zones after a threshold for the intensity was applied. A ratio between the intensity in each zone and the total intensity of the whole section was then calculated.

### Cell immunofluorescence and cell morphology analysis

The different cell subsets were FACS sorted in cold 5ml FACS tube containing 1ml of RPMI supplemented with 10% fetal calf serum, 1% L-glutamine (Gibco), 100 U ml−1 of penicillin– streptomycin, 1% nonessential amino acids, 1% sodium pyruvate and 0.05 mM β-mercaptoethanol. After washing in PBS, cells were resuspended in RPMI 1% FCS. 1.000 to 50.000 cells were allowed to adhere for 1 hour at 37°C to coverslips coated with 5 μg/cm^2^ poly-D-Lysine (Sigma). Cells were then washed in PBS 1x, fixed in 4% Paraformaldehyde (PFA) for 10 min, and permeabilized/blocked in PBS 1x containing 0.2% TritonX100, 2% FCS, 2% Rat serum, 2% Goat Serum and 2% Donkey serum. F-actin staining was performed at 4°C overnight in PBS 1x containing 0,1% TritonX100 2% FCS with Phalloidin AF405plus (Life technologies). Samples were washed in PBS 1x and mounted with Prolong Antifade Gold mounting medium (Life technologies). Images were acquired by spectral confocal microscope (Zeiss LSM 880) with 63x objective and analyzed with ImageJ. For the cell morphology, a binary image was created for each individual cell based on the F-actin staining. The circularity index (4π*area/perimeter^2) was then calculated with the adequate function of the software.

### Statistical analysis

All quantifications were performed with awareness of experimental groups, meaning not in a blinded fashion. Statistical parameters including the definitions and exact value of n (number of biological replicates and total number of experiments), and the types of the statistical tests are reported in the figures and corresponding figure legends. Statistical analyses were performed using Prism (GraphPad Software) or R statistical programming language. Statistical analysis was conducted on data with at least three biological replicates. Comparisons between groups were planned before statistical testing and target effect sizes were not predetermined. Error bars displayed on graphs represent the mean±SEM. Statistical significance was defined as * for P<0.05, ** for P<0.01, *** for P<0.001 and **** for P<0.0001.

## Acknowledgements

We thank CIPHE for the generation of *Siglech^iCre^* and *Pacsin1^LSL-tdT^* mice and their assistance in the breeding of mice, as well as the staff of the CIML mouse houses, flow cytometry, histology, genomics bioinformatics and imaging (ImagImm) core facilities. We thank S. Henri (CIML) for the generous gift of the *Zbtb46^GFP^* mice. We thank the two Master trainees, Alex Trinh and Dylan Anselmo, for technical assistance in some experiments. This research was funded by grants from the Fondation pour la Recherche Médicale (grants no. DEQ20110421284 and DEQ20180339172, Equipe Labellisée, to M.D.). We also thank the DCBIOL Labex (ANR-11-LABEX-0043, grant no. ANR-10-IDEX-0001-02 PSL*), the A*MIDEX project (grant no. ANR-11-IDEX-0001-02) funded by the French government’s Investissements d’Avenir program managed by the ANR, and institutional support from CNRS, INSERM, Aix-Marseille Université and Marseille Immunopole. This work was supported by the French National Research Agency through the Investments for the Future program (France-BioImaging, ANR-10-INBS-04).

## Author’s contributions

M.V. designed, performed and analysed the experiments, wrote the manuscript. N.C. performed part of the microscopy experiments, genotyping of mutant mice, generation and stabilization SCRIPT mice at triple homozygous state. T.-P.V.M. performed the bioinformatics analysis and contributed to write the manuscript. K.N. performed the crosses and genotyping for the generation and stabilization of the S-RFP and pDC-tdT mouse strains. G.B. contributed to the genotyping of mutant mice, and performed the crosses and genotyping for the generation and stabilization of the ZeST mice. L.G. trained M.V. for the generation of the FB5P-seq data. P.M. coordinated the generation and first level analysis of the FB5P-seq data. E.T. directed and contributed to fund the study, designed experiments, performed and analysed some experiments, and wrote the manuscript. M.D. directed and funded the study, designed experiments, contributed to the bioinformatics analysis, and wrote the manuscript.

---

**Figure S1.**
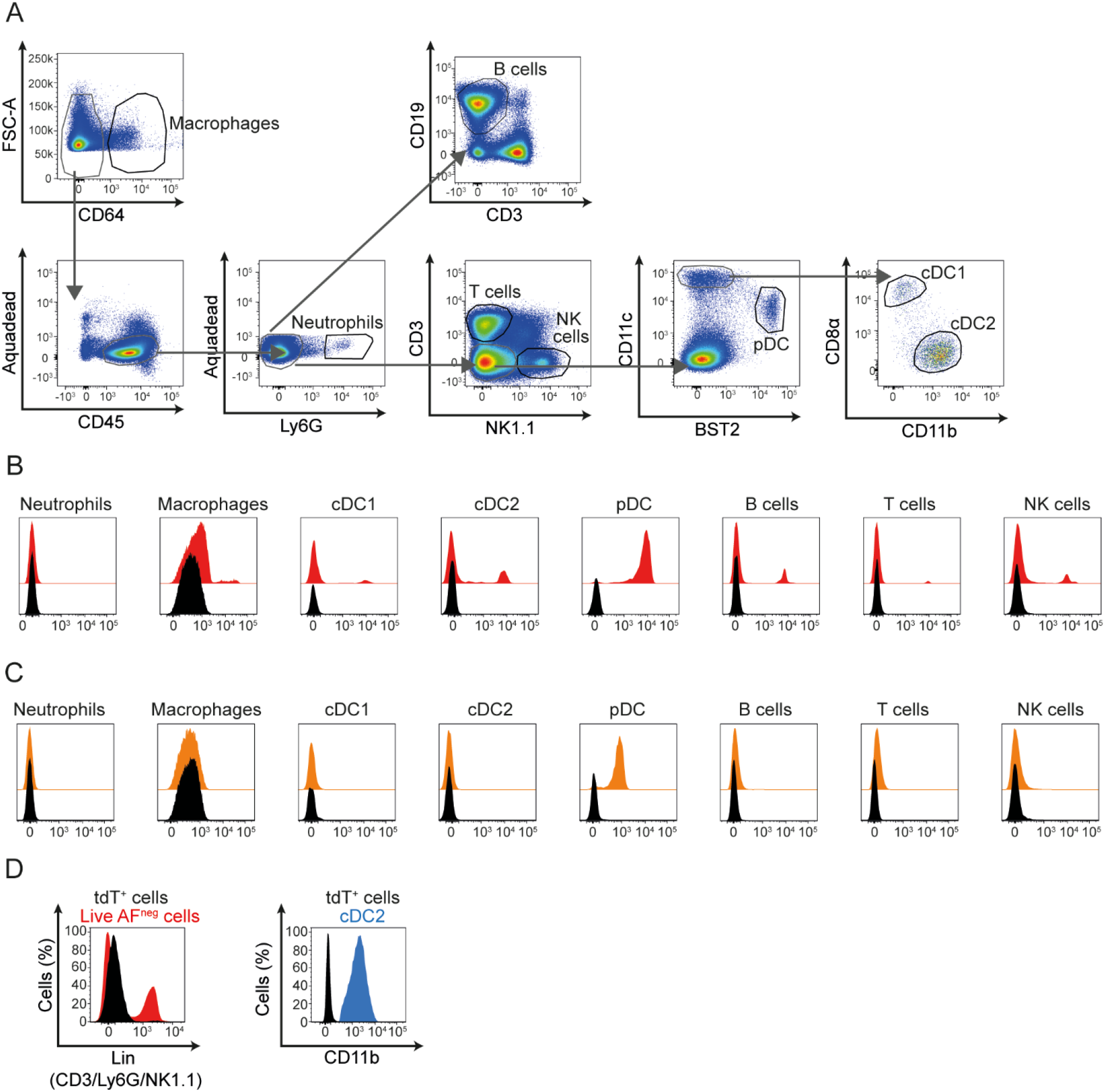
Expression of RFP and tdT in different immune lineages in S-RFP and pDC-Tomato mice. (A) Gating strategy used for the identification of the different immune cell types studied. (B-C) Splenocytes were isolated from S-RFP (B) or pDC-Tomato (C) animals, stained with fluorescently labeled antibodies and analyzed by flow cytometry. The expression of RFP (B, red histograms) or tdT (C, orange histograms) was evaluated in different immune cell types as indicated. (D) the expression of Lineage and CD11b was assessed on tdT^+^ cells and compared to Live non-autofluorescent cells (Lineage, left panel) or cDC2 (CD11b, right panel). Wt C57BL/6 mice were used as negative controls (black histograms). The data shown are from one mouse representative of 8 animals for (B) and 6 animals for (C-D) from 2 independent experiments.

**Figure S2.**
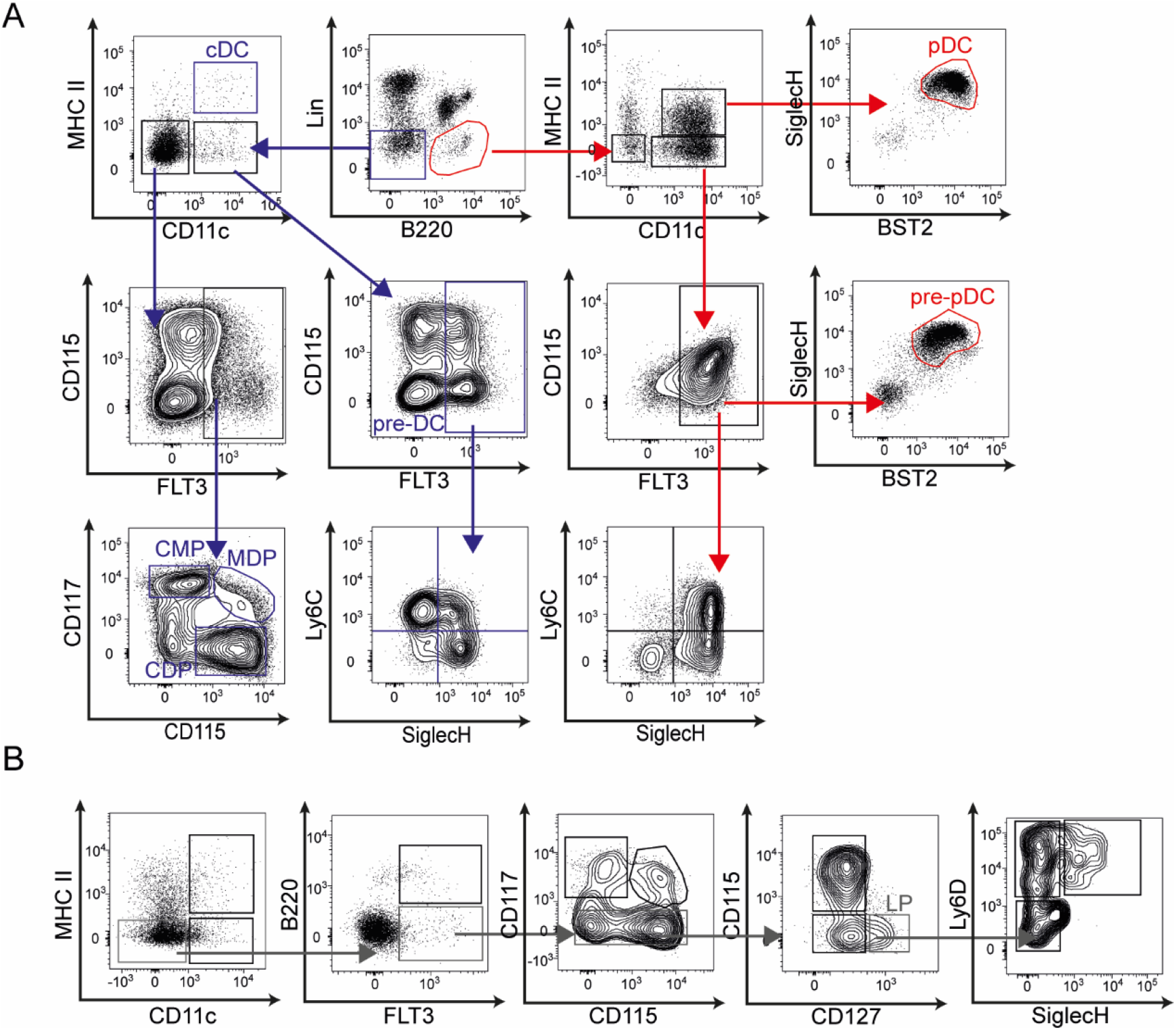
Gating strategy for bone marrow precursor analysis of pDC-Tomato mice. Bone marrow cells were isolated from pDC-Tomato animals, stained with fluorescently labeled antibodies and analyzed by flow cytometry. The gating strategy used to identify myeloid progenitors (A) or lymphoid progenitors (B) is depicted. The data shown are from one mouse representative of 5 pDC-Tomato animals from 2 independent experiments.

**Figure S3.**
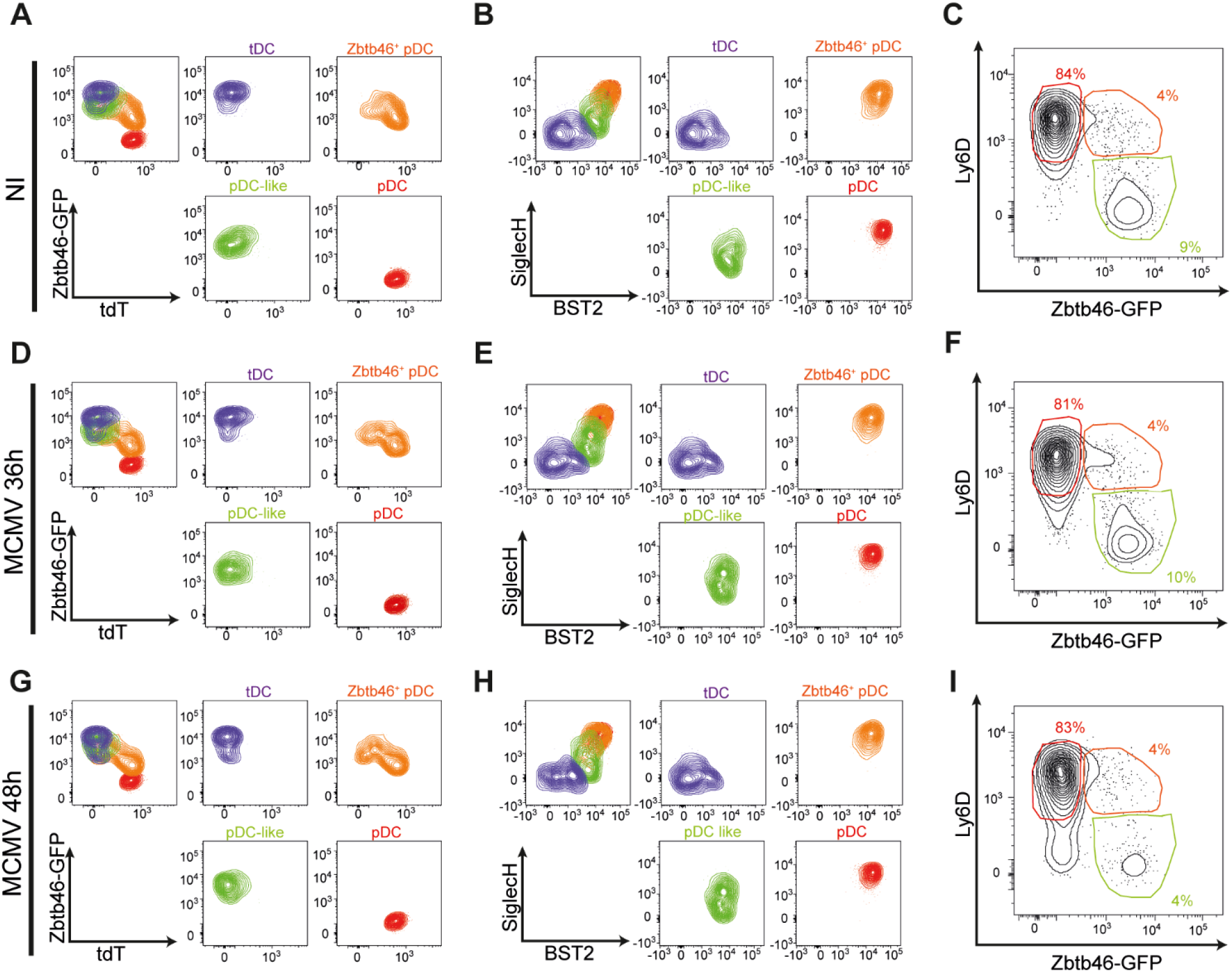
The phenotype and the proportions of Zbtb46^+^ pDC, tDC and pDC-like cells are relatively stable during MCMV infection. Splenocytes from ZeST mice were stained with fluorescently labeled antibodies and analyzed by flow cytometry. Representative contour plots from uninfected animals (A-C), MCMV-infected mice at 36h p.i (D-F) and from MCMV-infected mice at 48 p.i (G-I). The data shown are from one ZeST mouse representative of at least 10 uninfected animals (A-C), and for 7 MCMV-infected animals at 36h p.i (D-F) or 8 MCMV-infected animals at 48h p.i. (G-I).

**Figure S4.**
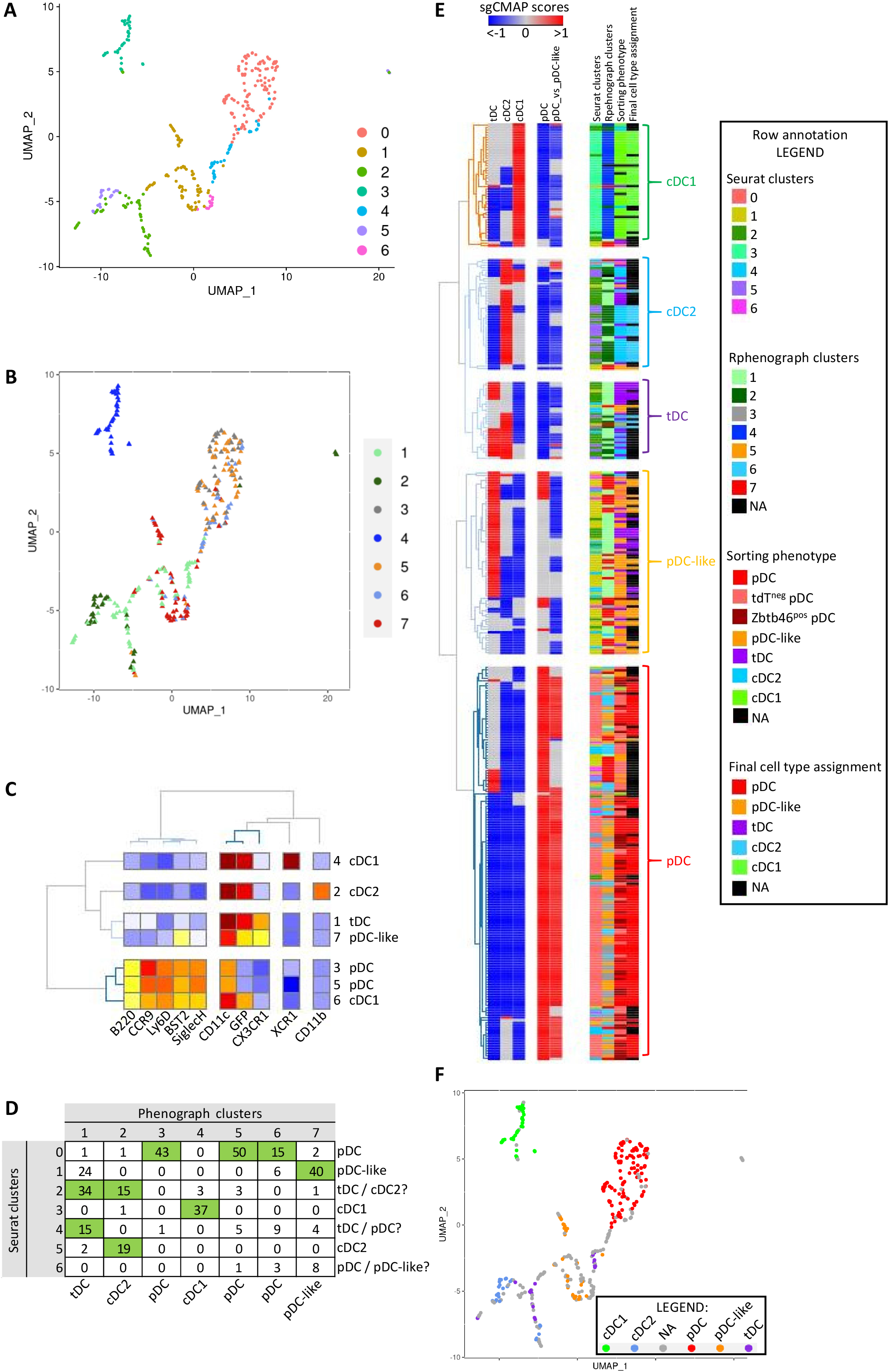
Analysis of the FB5P-seq data for cells from uninfected mice to generate DC type-specific signatures for helping annotation of the total FB5P-seq dataset. (A) Dimensionality reduction of the gene expression data performed using the UMAP algorithm with the Seurat pipeline for DC types isolated from the spleen of one uninfected ZeST (*Zbtb46^GFP^*;*Siglech^iCre^*;*Rosa26^LSL-tdT^*) mouse. Cells were index sorted into the 5 DC types studied using the gating strategy presented in Figure 3B, and used for single cell RNA sequencing using the FB5P-seq procedure. Cells that did not pass the FB5P-seq quality controls were removed from the analysis, leading to keeping 343 cells. The color code indicates belonging of the cells to the 7 Seurat clusters obtained. (B) Projection onto the UMAP space of the phenotype of cells based on their belonging to the R-Phenograph clusters (color code) obtained upon re-analysis of the fluorescent signals for 10 of the phenotypic markers acquired during index sorting as listed in panel C. (C) Annotation of the R-Phenograph clusters for DC type identity based on mean fluorescent intensities per marker and cluster as shown on the heatmap. (D) Matrix giving the number of cells belonging to the intersections between Seurat (rows) and R-Phenograph (column) clusters. Seurat clusters were annotated for cell types based on analysis of their marker genes. (E) Heatmap showing sgCMAP scores of individual cells (rows) for DC type-specific signatures (columns) generated from published independent RNA-seq datasets. Hierarchical clustering was performed, using the Pearson’s minus one metric for signatures (columns), and the Euclidian distance for individual cells (rows), with annotation of individual cells (rows) for i) belonging to Seurat clusters, ii) belonging to R-phenograph clusters, iii) sorting phenotype, and iv) final cell type assignment. Cells were assigned a cell type identity based on consistency between their belonging to Seurat and R-Phenograph clusters (green cells in matrix of panel D) and with their sgCMAP scores. This led to final cell type assignment for 205 cells, with 138 cells left non-annotated (NA). (F) Projection onto the UMAP space of the final cell type assignment.

**Figure S5.**
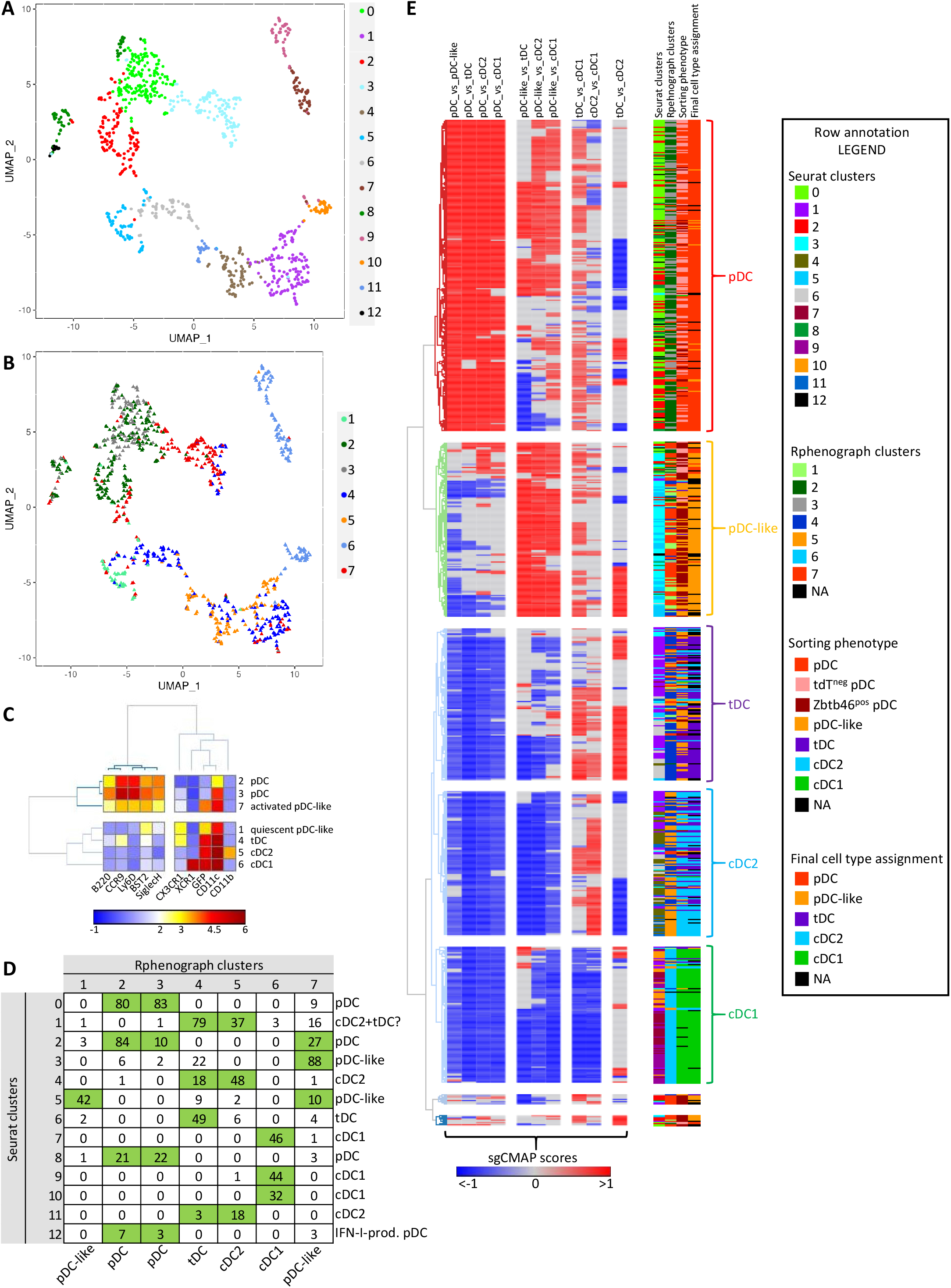
Analysis of the FB5P-seq data for all cells from uninfected and infected mice. A) Dimensionality reduction of the gene expression data performed using the UMAP algorithm with the Seurat pipeline for DC types isolated from the spleens of three ZeST (*Zbtb46^GFP^*;*Siglech^iCre^*;*Rosa26^LSL-tdT^*) mice (one uninfected, one MCMV-infected for 36h and the third infected for 48h). Cells were index sorted into the 5 DC types studied using the gating strategy presented in Figure 3B, and used for single cell RNA sequencing using the FB5P-seq procedure. Cells that did not pass the FB5P-seq quality controls or that turned out to be contaminants were removed from the analysis, leading to keeping 951 cells. The color code indicates belonging of the cells to the 13 Seurat clusters obtained. (B) Projection onto the UMAP space of the phenotype of cells based on their belonging to the R-Phenograph clusters (color code) obtained upon re-analysis of the fluorescent signals for 10 of the phenotypic markers acquired during index sorting as listed in panel C. (C) Annotation of the R-Phenograph clusters for DC type identity based on mean fluorescent intensities per marker and cluster as shown on the heatmap. (D) Matrix giving the number of cells belonging to the intersections between Seurat (rows) and R-Phenograp (column) clusters. Seurat clusters were annotated for cell types based on analysis of their marker genes. (E) Heatmap showing sgCMAP scores of individual cells (rows) for the DC type-specific sgCMAP signatures (columns) generated from the analysis of the data focused on cells from uninfected mice (see Figure S4). Hierarchical clustering was performed, using the Pearson’s minus one metric for signatures (columns), and the Euclidian distance for individual cells (rows), with annotation of individual cells (rows) for i) belonging to Seurat clusters, ii) belonging to R-phenograph clusters, iii) sorting phenotype, and iv) final cell type assignment. Cells were assigned a cell type identity based on consistency between their belonging to Seurat and R-Phenograph clusters (green cells in matrix of panel D), which was well corroborated by the sgCMAP scores. This led to final cell type assignment for 851 cells, with 100 cells left non-annotated (NA).

**Figure S6.**
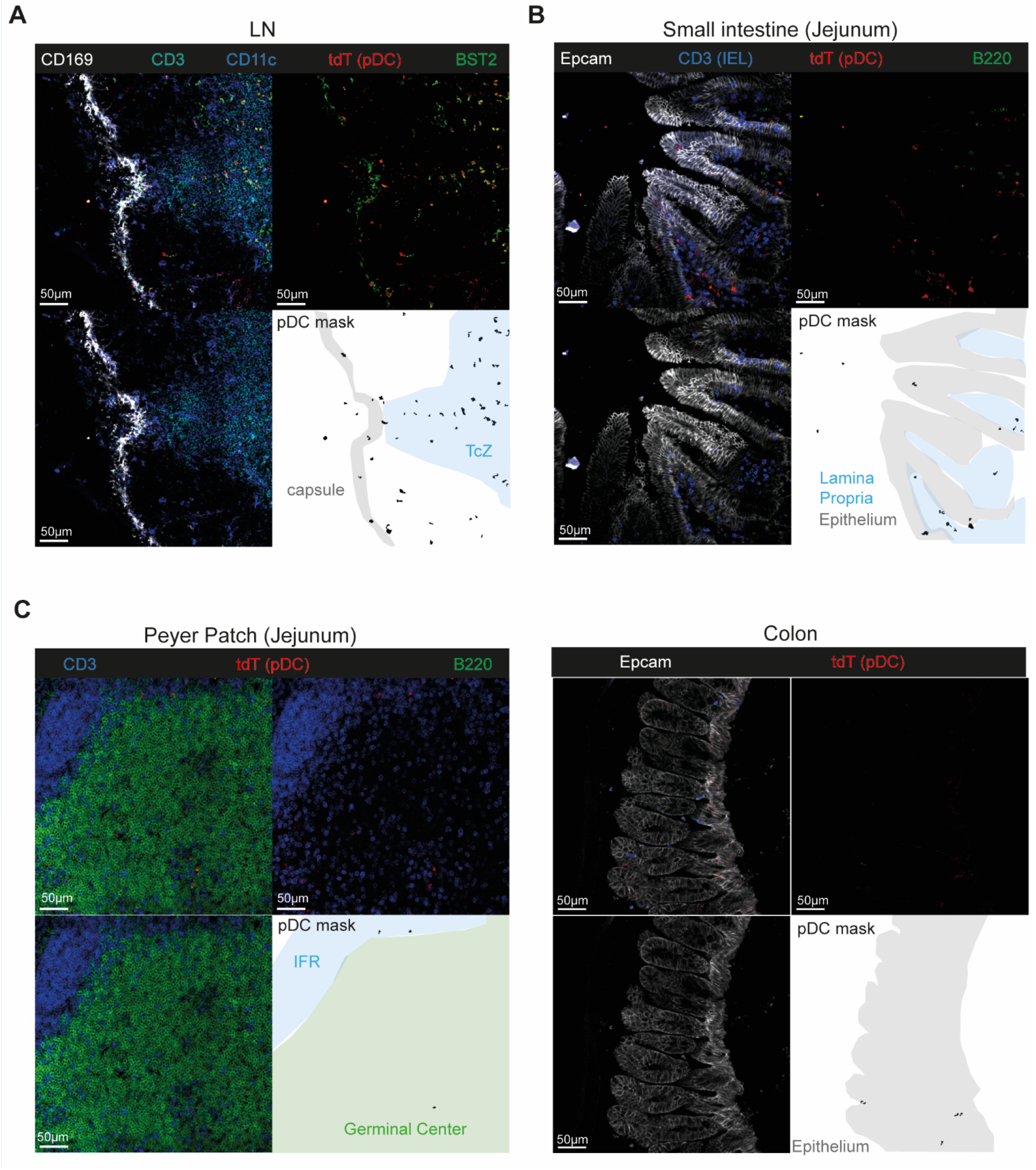
pDC-Tomato mice allow to determine pDC micro-anatomical location in the different organs. 20μm spleen cryosections from uninfected pDC-Tomato mice were stained with anti-tdT (red), anti-CD169 (white), anti-CD3 (cyan), anti-CD11c (blue) and anti-BST2 (green) antibodies (A) or with anti-tdT (red), anti-Epcam (white), anti-CD3 (IEL) and anti-B220 (green) antibodies (B-D). Representative images for LN of 3 mice, for small intestine of 4 mice and for colon of 3 mice.

**Figure S7.**
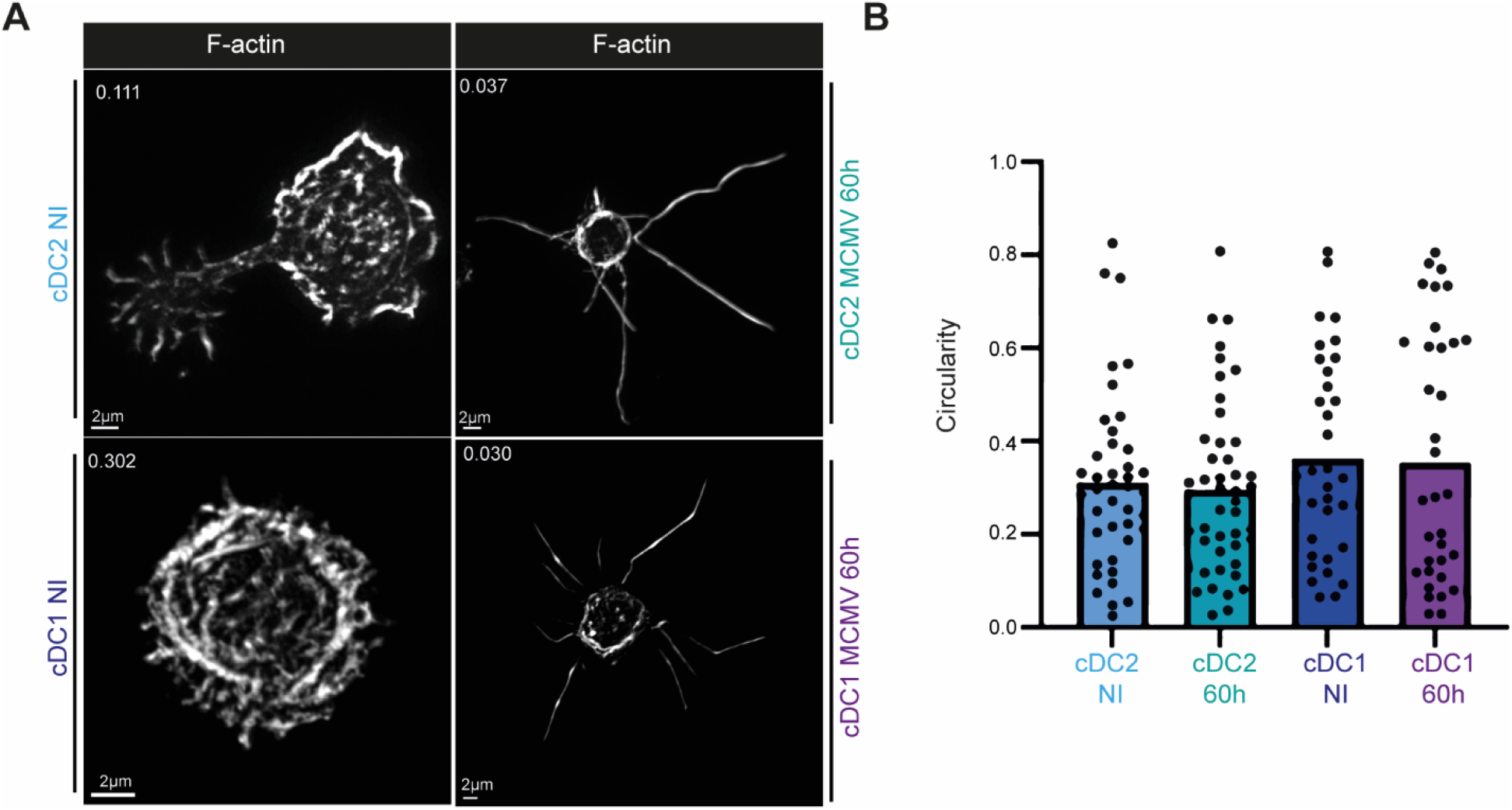
Circularity indices for cDC1 and cDC2 upon MCMV infection. (A-B) Quantitative and unbiased assessment of the cellular morphology of cDC1 and cDC2 isolated from 60h MCMV-infected SCRIPT mice, as compared to cDC1 and cDC2 from uninfected mice. (A) One representative confocal microscopy image of each DC type is shown. (B) The distribution of the circularity indices for individual cells across DC types is shown as dots, with the overlaid color bars showing the mean circularity indices of each DC type. The data shown are from 2 independent experiments, each performed with one mouse, with 35 to 46 individual cells analyzed for each DC type. The data are shown as mean±SEM.

**Figure S8.**
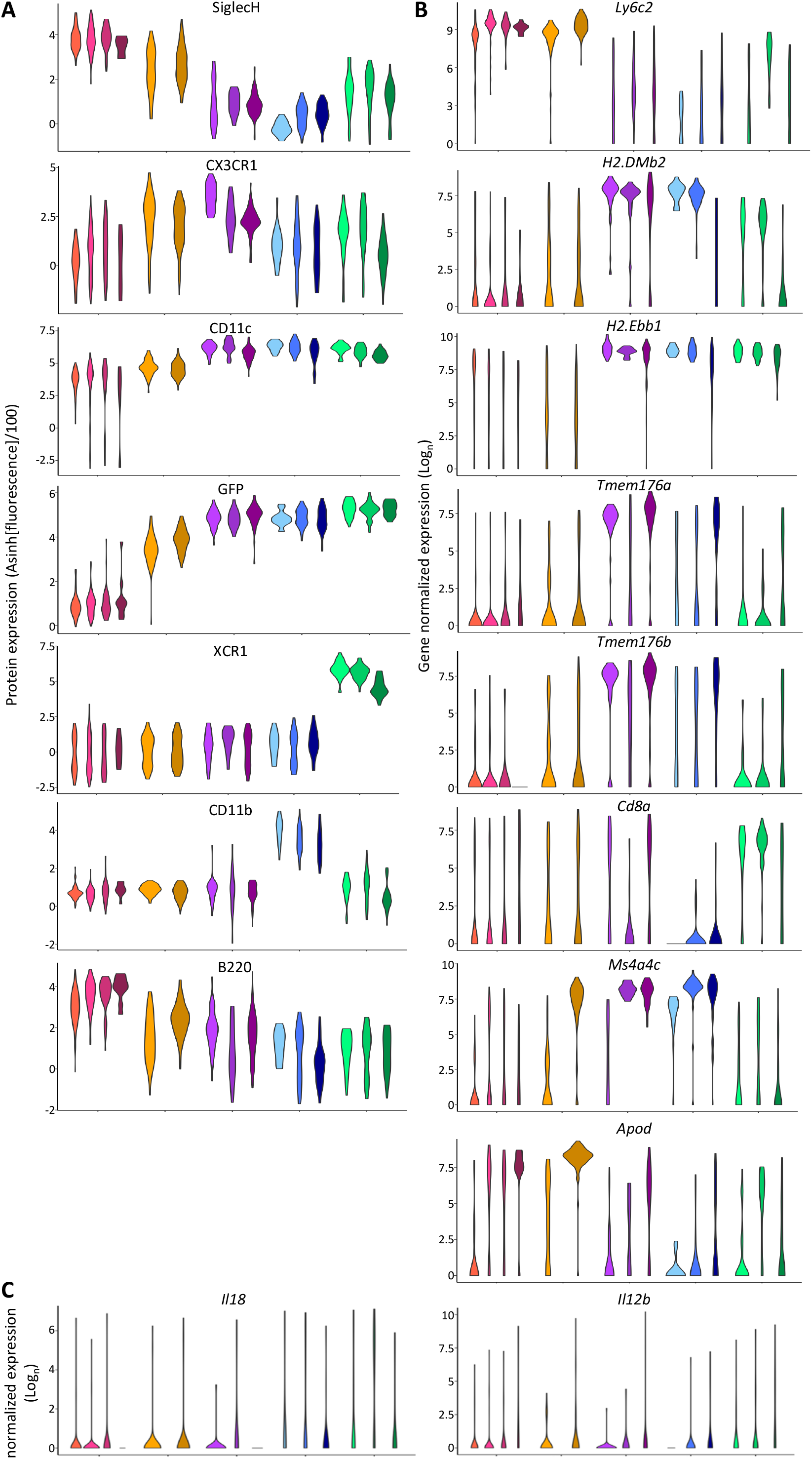
Expression of selected phenotypic markers and genes across DC types and activation states. (A) Violin plots showing the expression of phenotypic markers across DC types and activation states. (B-C) Violin plots showing mRNA expression levels of selected genes across DC types and activation states.

